# ERK signaling drives evolutionary expansion of the mammalian cerebral cortex

**DOI:** 10.1101/2023.08.08.552393

**Authors:** Mengge Sun, Yanjing Gao, Zhenmeiyu Li, Lin Yang, Guoping Liu, Zhejun Xu, Rongliang Guo, Yan You, Zhengang Yang

## Abstract

The molecular basis for cortical expansion during evolution remains largely unknown. Here, we found that FGF-extracellular signal-regulated kinase (ERK) signaling promotes the self-renewal and expansion of cortical radial glial (RG) cells.

Furthermore, FGF-ERK signaling induces *Bmp7* expression in cortical RG cells, which increases the length of the neurogenic period. We demonstrate that ERK signaling and SHH signaling mutually inhibit each other in cortical RG cells. We also provide evidence that ERK signaling is elevated in cortical RG cells during development and evolution. We conclude that the evolutionary expansion of the mammalian cortex, notably in human, is driven by ERK-BMP7-GLI3R signaling pathway in cortical RG cells, which participates in a positive feedback loop through antagonizing SHH signaling. We also propose that the molecular basis for cortical evolutionary dwarfism, exemplified by the lissencephalic mouse which originated from a larger and gyrencephalic ancestor, is due to mouse cortical RG cells receiving higher SHH signaling that antagonizes ERK signaling.

## INTRODUCTION

Insights into the evolution of the human cerebral cortex are beginning to come to light ^1–19^. Early human cortical development is driven by four types of neural stem cells which are the source for cortical glutaminergic pyramidal neurons (PyNs) and glia: ventricular zone (VZ) neuroepithelial cells, VZ full span radial glial (fRG) cells, VZ truncated radial glial (tRG) cells, and outer radial glial (oRG, also known as basal RG) cells which lie in the outer subventricular zone (outer SVZ, OSVZ) ^3, 5, 6, 8, 14, 20–26^. Human cortical neuroepithelial cells start to convert into fRG cells around gestational week 7 (GW7) and fRG cells give rise to oRG and tRG cells around GW16 ^24, 25, 27^. We recently provided evidence that oRG cells in the cortical OSVZ are mainly neurogenic, while tRG cells in the cortical VZ are mainly gliogenic, mainly giving rise to cortical astrocytes, oligodendrocytes and a subpopulation of olfactory bulb interneurons ^6, 25^. It seems that, the same limited repertoire of morphogens, such as BMPs, WNTs, FGFs, and SHH are used over and over again across ontogeny and phylogeny, and are involved in all steps of neural development from neural induction, and telencephalon patterning to neurogenesis and gliogenesis regulation ^1, 6, 28–31^.

Over the course of evolution, the primate cerebral cortex had a large increase in the number of neurons. Indeed, human cerebral cortex has the largest number of neurons (16.3 billion) ^32, 33^, which is a dominant contributor to the “seat” of intelligence and mind ^34^, whereas the elephant cerebral cortex, which has twice the mass of the human cerebral cortex, only has 5.6 billion neurons, which is also fewer than ∼8 billion neurons in the chimpanzee cerebral cortex ^35^.

In terms of cortical PyN numbers, there are mainly two determining factors. One is the size of cortical neuroepithelial and fRG cell pool present in the VZ at the beginning of cortical neurogenesis, which largely determines the PyN number and surface area of the cerebral cortex. This is known as “the radial unit hypothesis” ^3, 4^. Another is the length of the cortical neurogenic period. Indeed, humans have the longest cortical neurogenic period ^7^, allowing cortical RG cells to produce more PyNs ^2, 7, 9, 36^. Human cortical neurogenesis from fRG cells in the VZ extends from GW7 to GW16 ^24^, and cortical neurogenesis from oRG cells in the OSVZ extends from GW17 to GW26 ^6, 25^. We found that, during mammalian evolution (mouse, ferret, monkey, human), *BMP7* is expressed by increasing numbers of cortical RG cells ^6^. *BMP7* promotes neurogenesis, inhibits gliogenesis, and thereby increases the length of the neurogenic period, resulting in producing a large number of cortical PyNs. We demonstrate that BMP7 signaling in cortical RG cells inhibits SHH signaling through promoting GLI3 repressor (GLI3R) formation ^6^. Conversely, SHH signaling in cortical RG cells inhibits *BMP7* expression, but the molecular mechanisms governing this process remain poorly understood. We also do not know what kind of signals induces *BMP7* expression in cortical RG cells during development and evolution.

In the present study, we found that FGF-ERK signaling promotes cortical RG cell self-renewal and inhibits neural differentiation, which greatly expands the size of the founder population. Furthermore, ERK signaling induces *Bmp7* expression in cortical RG cells, which increases the length of the neurogenic period ^6^. We demonstrate that ERK signaling and SHH signaling mutually inhibit each other in cortical RG cells. We also provide evidence that ERK signaling is elevated in RG cells during cortical development and evolution. We conclude that the evolutionary expansion of the human cortex is driven by ERK-BMP7-GLI3R signaling pathway in cortical RG cells, which participates in a positive feedback loop through antagonizing the SHH signaling. In contrast, during mouse cortical development and evolution, we propose that relatively higher levels of SHH signaling antagonize ERK-BMP7 signaling in cortical RG cells, which reduces the size of cortical RG cell pool, represses *Bmp7* expression in dorsal RG cells, and thus reduces the length of the cortical neurogenic period. This provides a molecular basis for cortical evolutionary dwarfism, as exemplified by the lissencephalic mouse which originated from a larger and gyrencephalic ancestor ^7, 37, 38^. Taken together, we conclude that ERK signaling drives the evolutionary expansion of the mammalian cerebral cortex.

## RESULTS

### *Fgf8* overexpression promotes the expansion of cortical progenitors and induces *Bmp7* expression

Mouse cortical neurogenesis starts to take place following cortical patterning. During early cortical neurogenesis (i.e., E11.0-E12.5), *Fgf8, Fgf17, Fgf18, Spry1*, and *Spry2* are mainly expressed in the presumptive septum and extend into the medial cortex ^39–42^. Interestingly, *Bmp7* is also expressed in the medial cortex ^6, 43^. Furthermore, previous studies have shown that *Fgf8* and *Bmp7* are co-expressed in the mouse anterior neural ridge ^44^. These observations indicate that *Bmp7* expression might be induced by FGF signaling. To test this hypothesis, we overexpress *Fgf8* using *hGFAP-Cre* transgene mice to cross a conditionally overexpresses *Fgf8* line (*Rosa^Fgf8^* mice) (Figure S1A) ^45^. The hGFAP-Cre recombinase occurs specifically in cortical RG cells starting at E12.5 resulting in recombined floxed alleles in nearly all cortical RG cells and their progeny from E13.0 ^6, 46^. Consistent with our hypothesis, the strong *Bmp7* expression was observed in the E14.5 neocortical VZ of *hGFAP- Cre; Rosa^Fgf8^* mice, whereas *Bmp7* expression is restricted in the medial cortex of littermate controls (wild type, WT, or *hGFAP-Cre*) (Figure 1A). The interaction of FGF8 with FGFRs results in phosphorylation and nuclear translocation of ERK1/2, which phosphorylates target transcription factors (Figure S1B). We performed immunohistochemistry experiments using an antibody that detects phosphorylated forms of ERK1/2 (Corson et al. 2003). At E14.5, pERK^+^ cells in control mice are observed in the cortical VZ and SVZ, whereas we found a greatly enhanced expression of pERK in the cortex of *hGFAP-Cre; Rosa^Fgf8^* mice (Figure 1B). We also observed that FGF8 strongly induces SP8 expression (Figure 1C) ^47^. Notably, HOPX expression was significantly upregulated (Figure 1D). Constitutively overexpression of *Fgf8* also promotes the switch from cortical neurogenesis to gliogenesis, as EGFR and OLIG2 are strongly expressed in cortical VZ and SVZ in a lateral^high^-medial^low^ gradient (Figure 1E) ^48^, whereas their expression was absent in the control cortex at E14.5 (Figure 1E).

**Figure 1.**
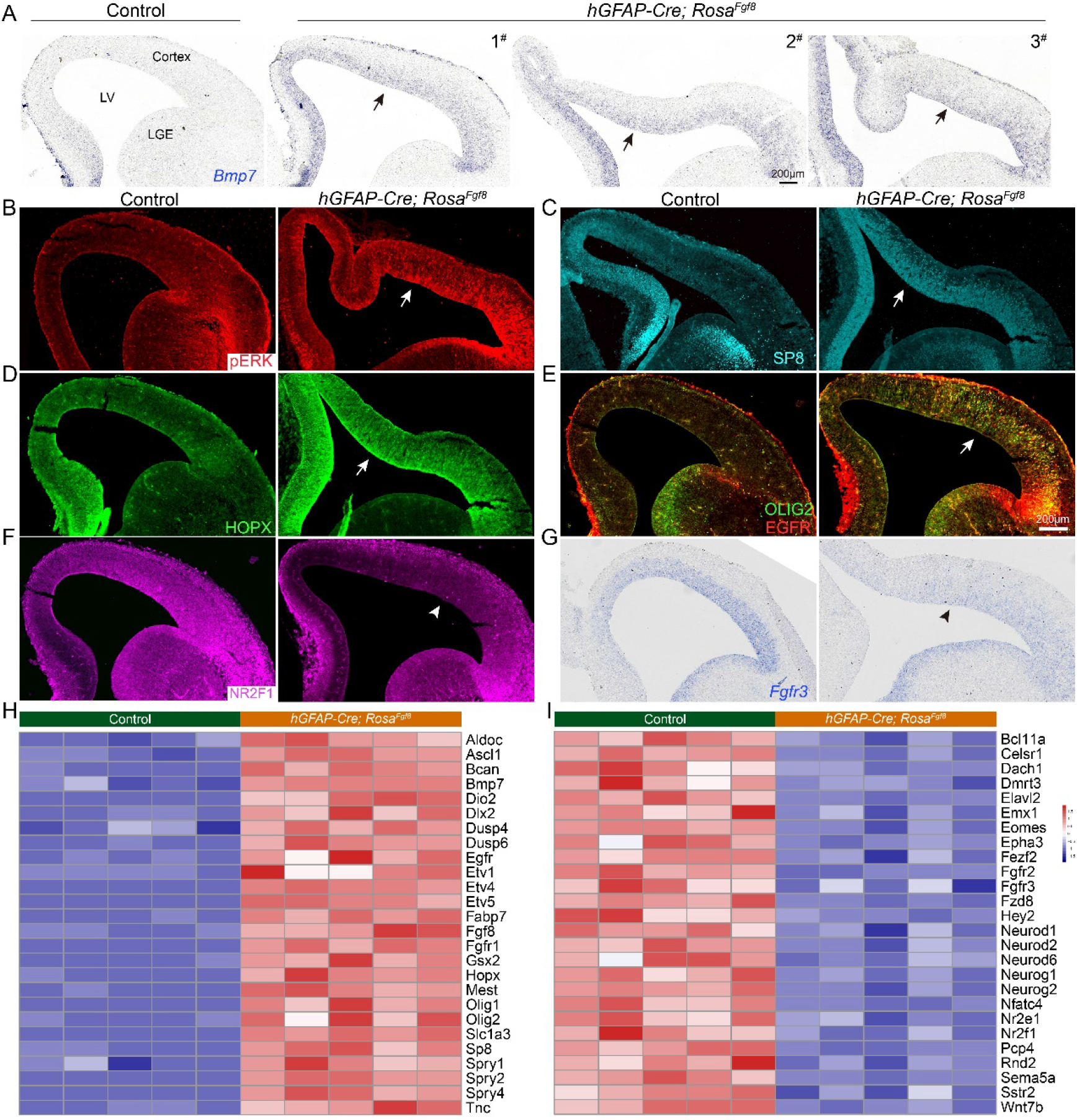
Overexpression of *Fgf8* expands the cortical progenitor pool and induces *Bmp7* expression in cortical RG cells. **(A)** mRNA in situ hybridizations on coronal sections through the telencephalon of E14.5 littermate control and *hGFAP- Cre; Rosa^Fgf8^* mouse embryos (n=3), showing significant upregulation of *Bmp7* expression in the cortex (arrows). Note that *Bmp7* expression is only detected in the medial cortex of littermate controls. LV, lateral ventricle. (B-G) Expression of pERK, SP8, HOPX, EGFR and OLIG2 was greatly increased (arrows), whereas expression of NR2F1 and *Fgfr3* was downregulated (arrowheads) in the cortex following *Fgf8* overexpression. (H, I) The heat map of bulk RNA-seq data showing expression of neurogenesis genes was downregulated, whereas expression of FGF8 signaling response genes, *Etv1, Etv4, Etv5, Mest, Sp8, Spry1, Spry2*, and *Spry4* was upregulated in the *hGFAP-Cre; Rosa^Fgf8^* cortex (n = 5) compared with littermate controls (n = 5) at E14.5.

FGF8 regulates cortical patterning by promoting rostral neocortical fates, and repressing caudal neocortical fates, in part through promoting *Etv1, Etv4, Etv5, Mest,* and *Sp8* expression, and repressing *Nr2f1* (*Coup-TFI*), *Emx2* and *Fgfr3* expression ^28, 39, 44, 47, 49–54^. hGFAP-Cre induction of ectopic *Fgf8* occurs about one day after neurogenesis has started, and about three days after the early stages in cortical regional patterning. None-the-less the ectopic *Fgf8* represses NR2F1 and *Fgfr3* expression in the E14.5 (Figure 1F, G), consistent with its known function ^28, 50, 52^. FGF signaling stimulates of cell proliferation ^55–58^. Accordingly, the *hGFAP-Cre; Rosa^Fgf8^* mice had increased expansion of the cortical VZ (Figure S2H).

To obtained unbiased transcriptional information on the effect of the ectopic *Fgf8*, we performed bulk RNA-Seq (Figure S1C). We compared gene expression profiles from the cortex of *hGFAP-Cre; Rosa^Fgf8^* mice and littermate controls at E14.5, and identified more than 4,000 genes that were either upregulated or downregulated (P < 0.05, Table S1). Selected differentially expressed genes are shown in the heat map (Figure 1I, J), which fully confirmed our immunostaining and mRNA in situ hybridization results. In addition, FGF signaling response genes *Etv1, Etv4, Etv5, Dusp4, Dusp6, Fgfr1, Spry1, Spry2, Spry4* and gliogenesis-related genes *Aldoc, Bcan, Dio2, Fabp7,* and *Tnc* were significantly upregulated, whereas neurogenesis-related genes *Bcl11a, Dmrt3, Emx1, Eomes* (*Tbr2*)*, Neurod1*, *Neurod2, Neurod6, Neurog1*, and *Neurog2* were downregulated (Figure 1I, J).

Taken together, our results reveal that overexpression of *Fgf8* promotes expansion of cortical progenitors and induces *Bmp7* expression in the cortical RG cells. However, sustained FGF8 signaling also promotes the cortical RG cell lineage progression and promotes the switch from cortical neurogenesis to gliogenesis.

### Sustained activation of ERK signaling promotes the expansion of cortical progenitors and induces *Bmp7* expression

FGF signaling is mediated by the activation of Ras/Raf/Mitogen-activated protein kinase/ERK kinase (MEK)/ERK cascade, phosphatidylinositol-4,5-bisphosphate 3- kinase (PI3K)-AKT, and Phospholipase C Gamma (PLCγ) (Figure S1B) ^57, 59^. ERK signaling can be activated in the *Rosa^MEK1DD^* mouse, which contains a Cre inducible constitutively active rat *Map2k1* and EGFP alleles. The mutant protein has 2 serine to aspartic acid substitutions (S218D/S222D) within the catalytic domain ^60, 61^.

We introduced the *hGFAP-Cre* allele into the *Rosa^MEK1DD^*mice, to study the effect of constitutively active *Map2k1* on intracellular signaling and gene expression. pERK is strongly upregulated in the cortical VZ and SVZ at E14.5 (Figure 2A). As expected, activated ERK induced *Bmp7* expression in the cortical VZ (Figure 2B). Furthermore, HOPX, ETV5, EGFR, and OLIG2 expression was also strongly upregulated (Figure 2C-F), whereas NR2F1 and *Fgfr3* expression was downregulated (Figure 2G, H). ERK signaling promotes progenitor cell proliferation ^62^. Accordingly, the VZ length was increased in the cortex of *hGFAP-Cre; Rosa^MEK1DD^* mice compared with littermate controls (Figure S2H).

**Figure 2.**
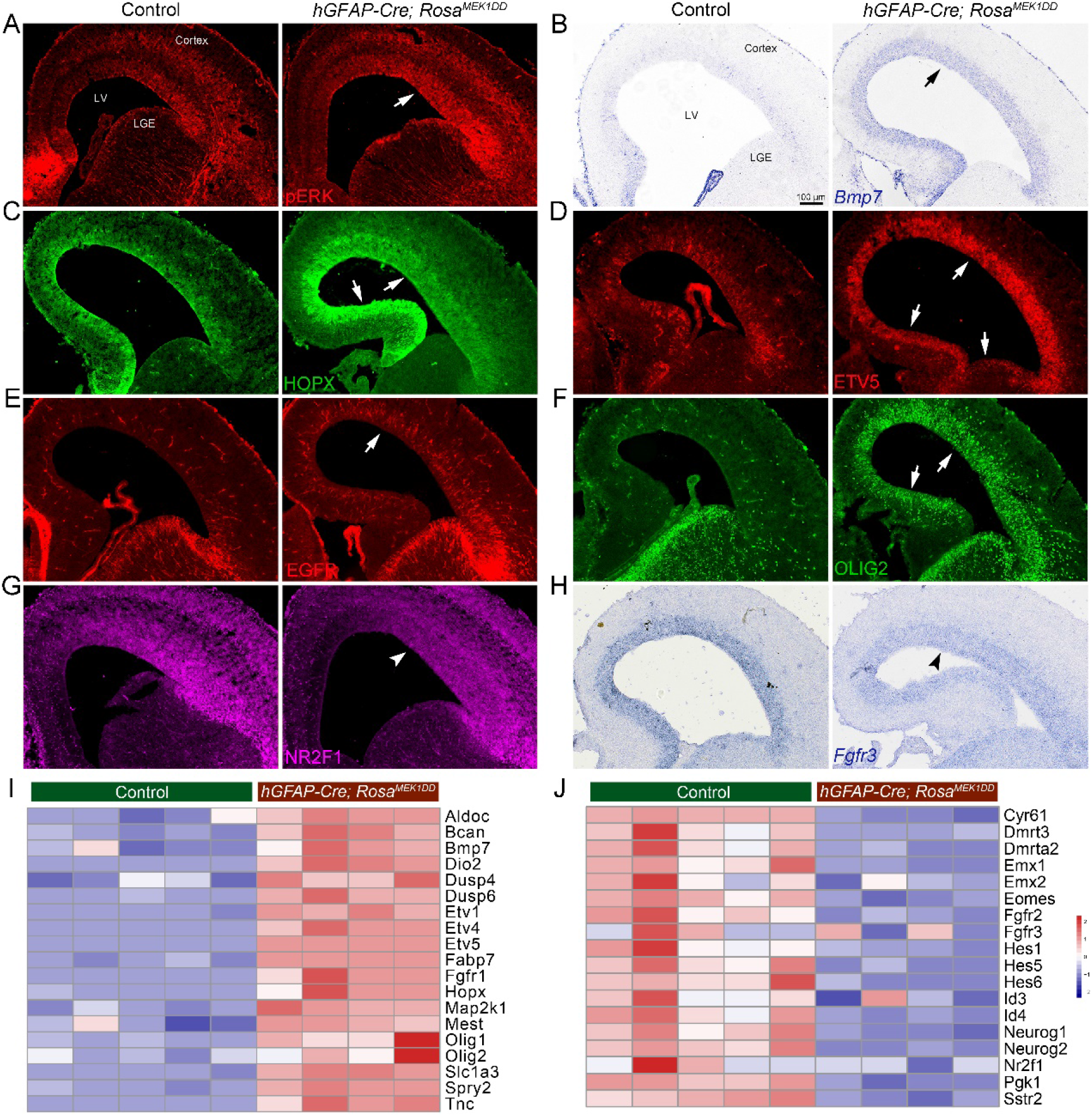
Constitutive activity of ERK signaling induces *Bmp7* expression in cortical RG cells. **(A-F)** Expression of pERK, *Bmp7*, HOPX, ETV5, EGFR, and OLIG2 was upregulated in the cortex of *hGFAP-Cre; Rosa^MEK1DD^* mice at E14.5 (arrows). (G, H) NR2F1 and *Fgfr3* expression was downregulated (arrowheads). (**I, J**) Heat map of selected differentially expressed genes in the E14.5 cortex of *hGFAP- Cre; Rosa^MEK1DD^* mice (n = 4) relatively to controls (n = 5).

Bulk RNA-Seq analysis confirmed the above results (Figure 2I, J, Table S2). There were more than 4, 000 genes that were either significantly upregulated or downregulated in the E14.5 cortex of *hGFAP-Cre; Rosa^MEK1DD^* mice (P < 0.05, Table S2). These misregulated genes were related to cell proliferation, survival, growth, metabolism, migration and differentiation, confirming that ERK signaling regulates these cellular processes in the developing cortex ^62^. Consistent with *Fgf8* overexpression, sustained ERK singling also promotes *Etv1, Etv4, Etv5, Dusp4, Dusp6, Spry2* expression, and promotes the onset of cortical gliogenesis (Figure 2I, J, Table S2).

Next, we examined *Nes-Cre; Rosa^MEK1DD^* mice at E14.5, and observed similar results to *hGFAP-Cre; Rosa^MEK1DD^* mice (Figure S2). Because Nes-Cre recombination occurs in the neural progenitors around E11.5, both in the pallium and subpallium ^63^, we found that *Bmp7* expression was also weakly upregulated in the lateral ganglionic eminence (LGE) at E14.5 (Figure S2A). This suggests that FGF-ERK signaling inducing *Bmp7* expression in RG cells might be a common rule in certain brain regions, including the neocortex and LGE. In summary, increase in FGF8-ERK signaling promotes the expansion of cortical progenitors and induces *Bmp7* expression in cortical RG cells, and sustained FGF8-ERK activity promotes the switch from the cortical neurogenesis to gliogenesis.

### Loss of cortical ERK signaling results in loss of *Bmp7* expression, depletion of cortical RG cells and premature neural differentiation

To further examine the function of ERK signaling in cortical RG cells, we conditionally deleted floxed alleles of core components of ERK pathway: *Map2k1* and *Map2k2* (using *Emx1-Cre* line; Figure S1A, B) ^64^. We have found that *Map2k1* and *Map2k2* double deletion (*Emx1-Cre; Map2k1/2-dcko*) depleted expression of pERK and HOPX at E14.5 (Figure 3A, B). Loss of ERK signaling also depleted the expression of *Bmp7* in the medial cortex (Figure 3C). On the other hand, expression of NR2F1 and *Fgfr3* was greatly increased, especially in the medial cortex (Figure 3D, E). We observed that a subset of EOMES^+^ PyN IPCs was located at the VZ surface in *Emx1-Cre; Map2k1/2-dcko* mice, while in controls, most EOMES^+^ PyN IPCs were located in the SVZ (Figure 3F), indicative of premature differentiation into PyN IPCs. Quantification of the E14.5 cortical VZ length showed a statistically significant decrease of VZ extension (Figure S2H), suggesting a reduction of cortical RG cells in *Emx1-Cre; Map2k1/2-dcko* mice.

**Figure 3.**
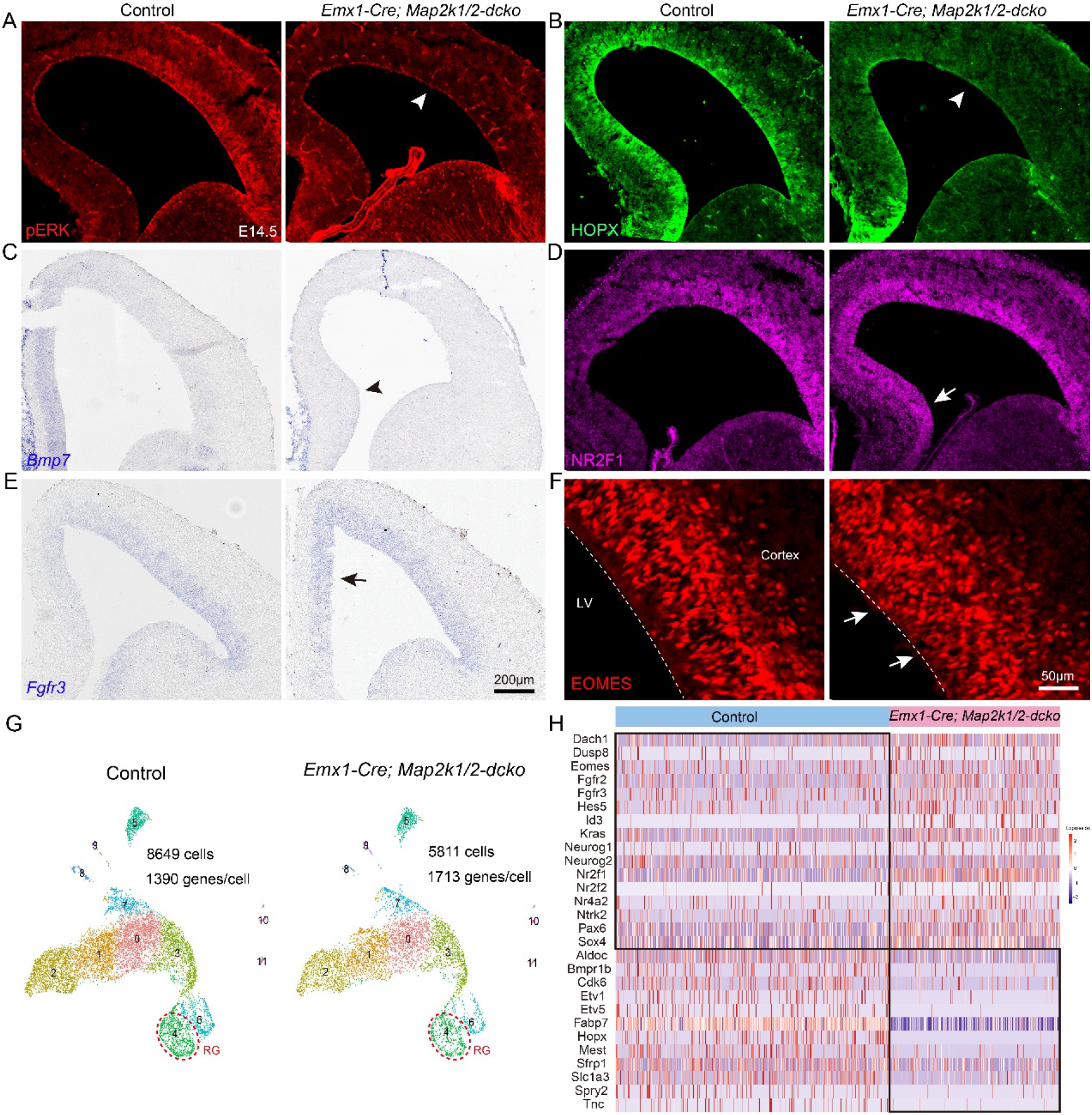
Deletion of *Map2k1* and *Map2k2* genes in the developing cortex leads to loss of ERK signaling and premature neural differentiation. **(A-C)** Expression of pERK, HOPX, and *Bmp7* was lost specifically in the cortex of *Emx1-Cre; Map2k1/2-dcko* mice at E14.5 (arrowheads). (D, E) NR2F1 and *Fgfr3* expression was increased in the cortex, especially in the medial cortex (arrows). (F) Immunostaining of EOMES, a marker for PyN IPCs, showing that PyN IPCs located in the VZ surface in *Emx1-Cre; Map2k1/2-dcko* mice at E14.5 (arrows). (G, H) scRNA-Seq analysis and heat map showing expression of selected ERK signaling response genes in cortical RG cells of *Emx1-Cre; Map2k1/2-dcko* mice relative to littermate controls at E14.5. Note that the expression of *Etv1, Etv5, Hopx, Spry2*, and *Tnc* was completely lost in cortical RG cells in *Emx1-Cre; Map2k1/2-dcko* mice.

At E17.0, expression of pERK and HOPX was still completely absent in the cortex of *Emx1-Cre; Map2k1/2-dcko* mice (Figure S3A, B); the thickness of the cortical VZ and SVZ was significantly reduced (Figure S3C-E). Remarkably, the number of cortical PAX6^+^ cells and EOMES^+^ cells was reduced to 50% lower in *Emx1-Cre; Map2k1/2- dcko* mice compared with controls (Figure S3C-G). These observations are consistent with previous results by deletion of all three *Fgf receptor* genes ^65^ or deletion of both *Mapk1 (Erk2)* and *Mapk3 (Erk1)* genes ^66^, indicating that FGF-ERK signaling promotes the self-renew of cortical RG cell and represses neural differentiation.

scRNA-Seq analysis of the E14.5 cortex of *Emx1-Cre; Map2k1/2-dcko* mice and littermate controls confirmed the above results (Figure 3G, H, Table S3). Without ERK signaling, expression of *Eomes, Fgfr2, Fgfr3, Hes5, Neurog1, Nr2f1*, and *Nr2f2* in cortical RG cells was upregulated, indicating premature neural differentiation, whereas expression of ERK response genes *Aldoc, Cdk6, Fabp7*, and *Slc1a3* was significantly downregulated, and expression of *Etv1, Etv5, Hopx, Mest, Spry2* and *Tnc* in cortical RG cells was nearly completely lost (Figure 3H).

In summary, our results reveal that, ERK signaling is required for maintaining the self-renewal and undifferentiated state of cortical RG cells; ERK signaling is absolutely required for the cortical expression of *Bmp7* and HOPX, and ERK signaling represses cortical expression of NR2F1 and *Fgfr3*. Previously, we have shown that *Bmp7* overexpression in the cortex increases HOPX expression ^6^, suggesting that both ERK and BMP7 signaling promote HOPX expression.

### BMP7-SMAD signaling promotes GLI3R formation

BMP7 binding to the type I and type II BMP receptors on the cell membrane results in the phosphorylation and activation of SMAD1/5/9 proteins, which form a complex with SMAD4 that together regulates gene expression ^67^. This is the BMP canonical Smad pathway ^67, 68^. When *Bmp7* was delivered into the cortical VZ by in utero electroporation (IUE) of *pCAG-Bmp7-Gfp* at E14.5 or E15.5, Western blotting showed increased ratio of GLI3R/GLI3FL (repressor/full-length forms of GLI3) in the E16.5 or E17.5 cortex, mainly due to excess production of GLI3R, which antagonizes SHH signaling ^6^. Expression of a constitutively active SMO protein that drives SHH signaling (*hGFAP-Cre; Rosa^SmoM2^*mice) significantly decreased GLI3R expression and increased ERFR expression ^6^. Here we overexpressed *Bmp7* in the cortical VZ of mice with genetic enhancement in SHH signaling and genetic defect in SMAD signaling (*hGFAP-Cre; Rosa^SmoM2^*mice and *hGFAP-Cre; Rosa^SmoM2^; Smad4^F/F^*) at E15.0 by IUE of *pCAG-Bmp7-Gfp*. Analysis at E17.0 in the *hGFAP-Cre; Rosa^SmoM2^ mice* showed that BMP7 induced increased expression of ID3 and pSMAD1/5/9, and decreased expression of EGFR, *Gli1*, *Ptch1*, and *Gas1* (Figure 4A-H). Importantly, the *Bmp7*-IUE cortex of *hGFAP-Cre; Rosa^SmoM2^; Smad4^F/F^* mice showed none of these gene expression was changes (Figure 4C-H). These results further demonstrate that BMP7 inhibits EGFR expression and promotes GLI3R formation through the canonical SMAD pathway.

**Figure 4.**
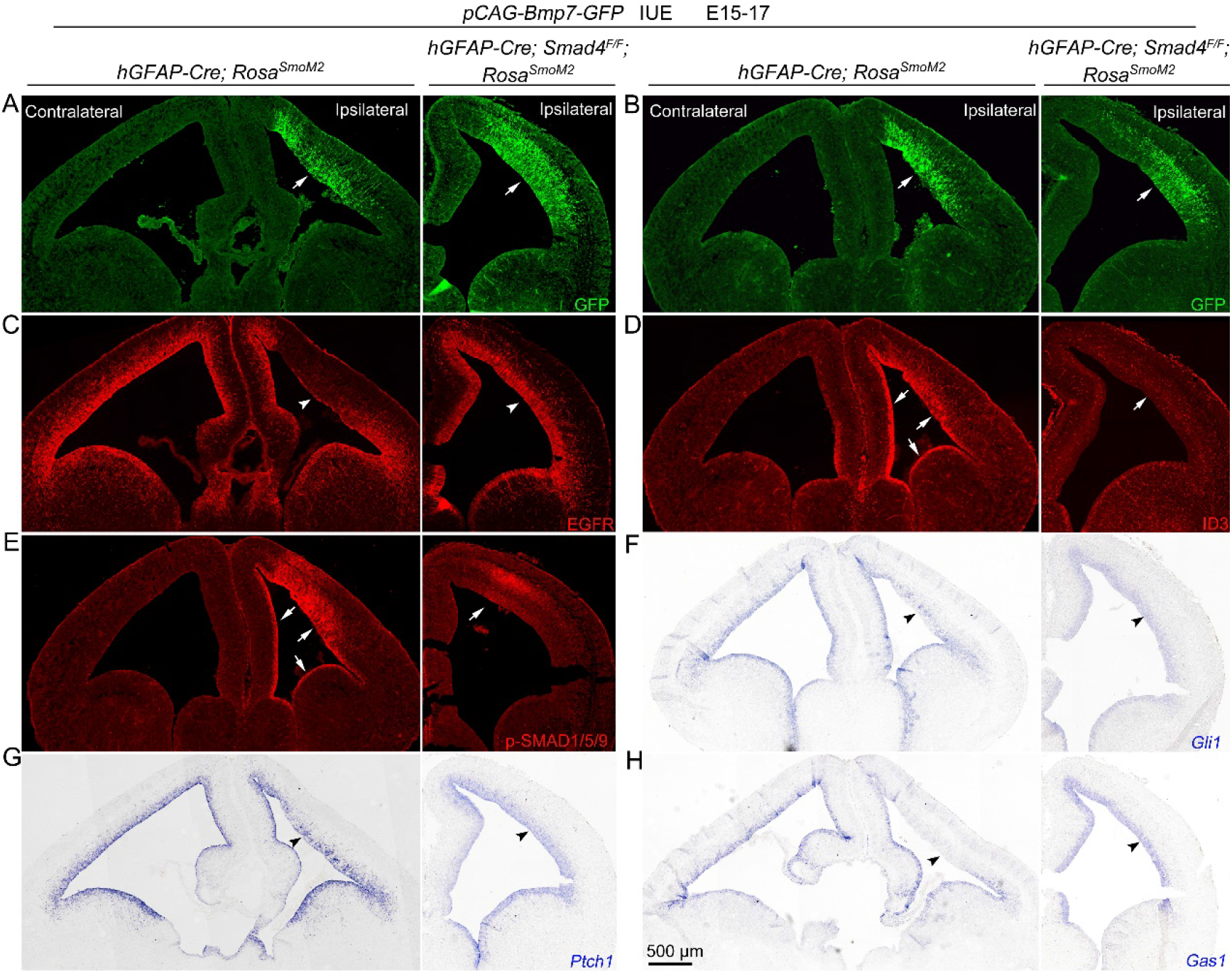
BMP7 promotes GLI3R generation in cortical RG cells through canonical SMAD pathway. **(A-H)** Overexpression of *pCAG-Bmp7-Gfp* in the E15.0 cortex by IUE induces ID3 and pSMAD1/5/9 expression (arrows), and represses EGFR, *Gli1*, *Ptch1*, and *Gas1* expression (arrowheads) in the E17.0 cortex of *hGFAP-Cre; Rosa^SmoM2^* mice, but not in *hGFAP-Cre; Rosa^SmoM2^; Smad4^F/F^* mice. Note the widespread upregulation of ID3 (D) and pSMAD1/5/9 (E) in the VZ/SVZ of the cortex, LGE and septum after cortical *Bmp7*-IUE. This is likely due to ultrasensitivity of BMP receptors, as the expression level of *Bmp7* is very low in the medial cortex of *hGFAP-Cre; Rosa^SmoM2^* mice. This also suggests that BMP7 is a diffusible morphogen, and may diffuse through both the extracellular space and the CSF.

### SHH-SMO signaling antagonizes ERK signaling in mouse cortical RG cells

In general, SHH signaling (SHH concentration) exhibits in a ventral^high^-dorsal^low^ and caudal^high^-rostral^low^ gradient, whereas FGF-ERK signaling exhibits in a rostral^high^- caudal^low^ gradient during telencephalon patterning and development ^30, 69, 70^. SHH and FGF8 have a reciprocal regulatory relationship during telencephalon patterning. SHH is required for maintaining *Fgf8* expression ^71^, and FGF8 is required for *Shh* induction ^28^. We examined SHH-SMO function in mouse cortical RG cells after neurogenesis has started bypassing the cortical patterning stage. In the E14.5 cortex of *hGFAP- Cre; Rosa^SmoM2^* mice, the expression of *Gli1, Fgf15, Fgfr3*, NR2F1, EGFR and OLIG2 was significantly upregulated, whereas the expression of pERK and HOPX was greatly reduced in RG cells compared with littermate controls (Figure 5A-H). These observations were further confirmed by scRNA-Seq analysis (Figure 5I, J, Table S4), suggesting that SHH-SMO signaling antagonizes ERK signaling in cortical RG cells after the cortical patterning stage, and sustained SHH-SMO activity promotes cortical gliogenesis ^72–74^.

**Figure 5.**
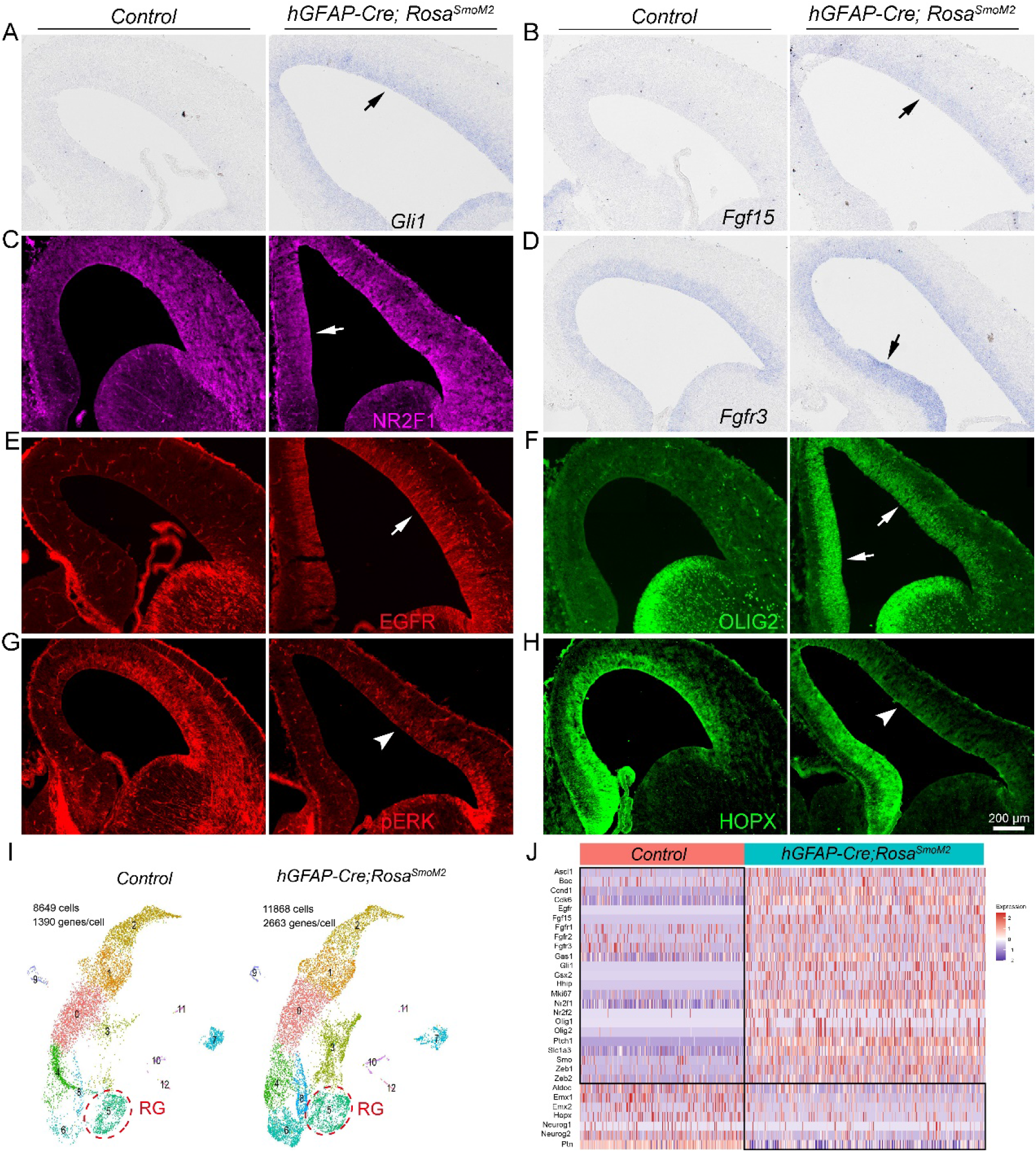
The SHH-SMO signaling antagonizes ERK signaling in cortical RG cells. **(A-H)** Expression of *Gli1*, *Fgf15*, NR2F1, *Fgfr3*, EGFR, and OLIG2 was greatly upregulated (arrows), whereas expression of pERK and HOPX was severely reduced (arrowheads) in the cortex of *hGFAP-Cre; Rosa^SmoM2^*mice compared with littermate controls at E14.5. (I, J) scRNA-Seq analysis and heat map showing expression of key signature genes in cortical RG cells of *hGFAP-Cre; Rosa^SmoM2^* mice relative to controls at E14.5.

We next tested whether ERK signaling is increased in cortical RG cells when SHH-SMO function is absent, using *hGFAP-Cre; Smo^F/F^* mice. At E17.0, expression of pERK, HOPX and *Bmp7* was upregulated the cortical VZ (Figure S4B-D) ^6^. scRNA-Seq analysis of cortical RG cells in *hGFAP-Cre; Smo^F/F^*mice and littermate *Smo^F/F^* control mice at E18.0 revealed that expression of *Aldoc, Bmp7, Dio2, Etv1, Etv4, Etv5, Fgfr2, Gfap, Hopx, Nog,* and *Tnc* was significantly upregulated (Figure S1D, S4I) ^6^. These results support the idea that, without SHH-SMO function, ERK signaling in cortical RG cells is enhanced.

SHH-SMO signaling inhibits *Bmp7* expression in the mouse cortex ^6^, most likely due to its antagonizing ERK signaling. To test this further, we reduced FGF-ERK signaling by IUE a soluble form of *FGFR3c* (*sFGFR3c*) in the E14.0 cortex of *hGFAP-Cre; Smo^F/F^* mice (Figure S4A, E). sFGFR3c is a high-affinity FGF receptor isoform that sequesters endogenous FGF family members, blocking their ability to activate endogenous receptors ^49, 75^. Both in E17.0 control and *hGFAP-Cre; Smo^F/F^* mice, the *sFGFR3*-IUE cortex showed decreased expression of pERK and HOPX (Figure S4F, G). Furthermore, *Bmp7* upregulation disappeared upon *sFGFR3c* overexpression (Figure S4H), suggesting that an increase in ERK signaling is crucial to induce *Bmp7* expression in the dorsal cortex of *hGFAP-Cre; Smo^F/F^* mice.

Taken together, our analysis of SHH-SMO gain of function and loss of function phenotypes in mice demonstrates that SHH signaling antagonizes ERK signaling in cortical RG cells after cortical patterning; this then contributes to preventing the spread of *Bmp7* expression from the medial cortex to the dorsal cortex.

### ERK signaling activity is evolutionarily elevated in mammalian cortical RG cells

We recently found that *BMP7* is expressed by increasing numbers of cortical RG cells during mammalian evolution (mouse, ferret, monkey, and human) ^6^. Because *BMP7* expression in RG cells is dependent on ERK signaling, we suspect that ERK signaling might be also elevated in mammalian cortical RG cells during evolution. We first examined the expression of ERK signaling response genes in cortical neurogenic fRG cells of mouse at E15.5 (whole cortex) ^76^, ferret at E39 (whole cortex) ^6^, rhesus monkey at E78 (visual cortex) ^77^, and human at GW14 (prefrontal cortex) ^78^ by comparative scRNA-Seq analyses (Figure S5A) ^6^, and found that, in general, there is an increase in the expression of *BMP7*, *ETV1, ETV5, FGF2, HOPX, SLC1A2, SLC1A3,* and *SPRY2* in cortical fRG cells during evolution (Figure S5B-D). The most prominent feature is the elevated expression of *SPRY1* in ferret, monkey and human cortical fRG cells (Figure S5D), whereas very few *SPRY1*-expressing fRG cells are in the mouse dorsal cortex ^40^. The expression of *SPRY* is dependent on ERK signaling, and SPRY participates in the negative-feedback control of ERK signaling ^40, 79^. We suspect that because mouse cortical ERK signaling is relatively weak ^80^, it fails to induce *Spry1* expression in dorsal cortical RG cells.

We next examined the expression of ERK signaling response genes in human cortical fRG and oRG cells (they are both neurogenic) at GW12, GW14, GW18, GW22, GW23 and GW26 by analyzing published scRNA-Seq dataset (Figure S6A) ^6, 78, 81^. As human development proceeds, more cortical RG cells express *ALDOC, ACSBG1*, *BMP7, DIO2, ERG1, ETV1, FGF1, FGF2, GFAP, FGFR1, FGFR2, HOPX, SLC1A2, SLC1A3, SPRY1, and SPRY2* (Figure S6B-E), indicating an elevation of ERK signaling in human cortical fRG and oRG cells with increasing gestational age.

To validate above scRNA-Seq analysis results, we performed immunostaining and mRNA in situ hybridization experiments to examine expression of pERK, HOPX and *BMP7* in the E33 ferret cortex, a developmental time point that is comparable to the E14.5 mouse cortex, as the cortical plate is newly formed in both animals at this stage (Figure S7A, B) ^82–84^. In the E14.5 mouse cortex, *Bmp7* mRNA was only detected in the medial cortex (Figure S7E); pERK and HOPX were main expressed in the whole cortical VZ, albeit relatively weak (Figure S7C, D). On the other hand, in the E33 ferret cortex, strong pERK and HOPX expression was seen in cortical fRG cells (Figure S7F, G). Furthermore, *Bmp7* mRNA expression extends much more broadly, from the medial cortex to dorsal cortex (Figure S7H), providing evidence that relatively stronger ERK signaling is present in the ferret cortical VZ than that in the mouse cortical VZ.

We also examined pERK and HOPX expression in the human cortical oRG and tRG cells at GW18. scRNA-Seq analysis of the GW18 human cortex from published datasets ^81^, revealed that human cortical oRG cells are neurogenic, while tRG cells are gliogenic ^6^, and ERK signaling response genes *ACSBG1, BMP7, ETV1, ETV5, FGFR1, GFAP, HOPX, S100B, SLC1A2, SLC1A3, SPRY1, SPRY2, and TNC* were highly expressed in oRG cells than that in tRG cells (Figure S8A-C, Table S5). Consistent with this bioinformatic analysis, we observed stronger pERK immunoreactivity in the soma and radial fibers of oRG cells in the OSVZ compared with tRG cells in the cortical VZ (Figure 7A-D), and most HOPX^+^ RG cells were co-expressed with pERK (Figure 7C, D). We propose that, similar to the mouse cortex, cortical expression of *BMP7* and HOPX in other mammals, including humans, are also induced by ERK signaling.

**Figure 6.**
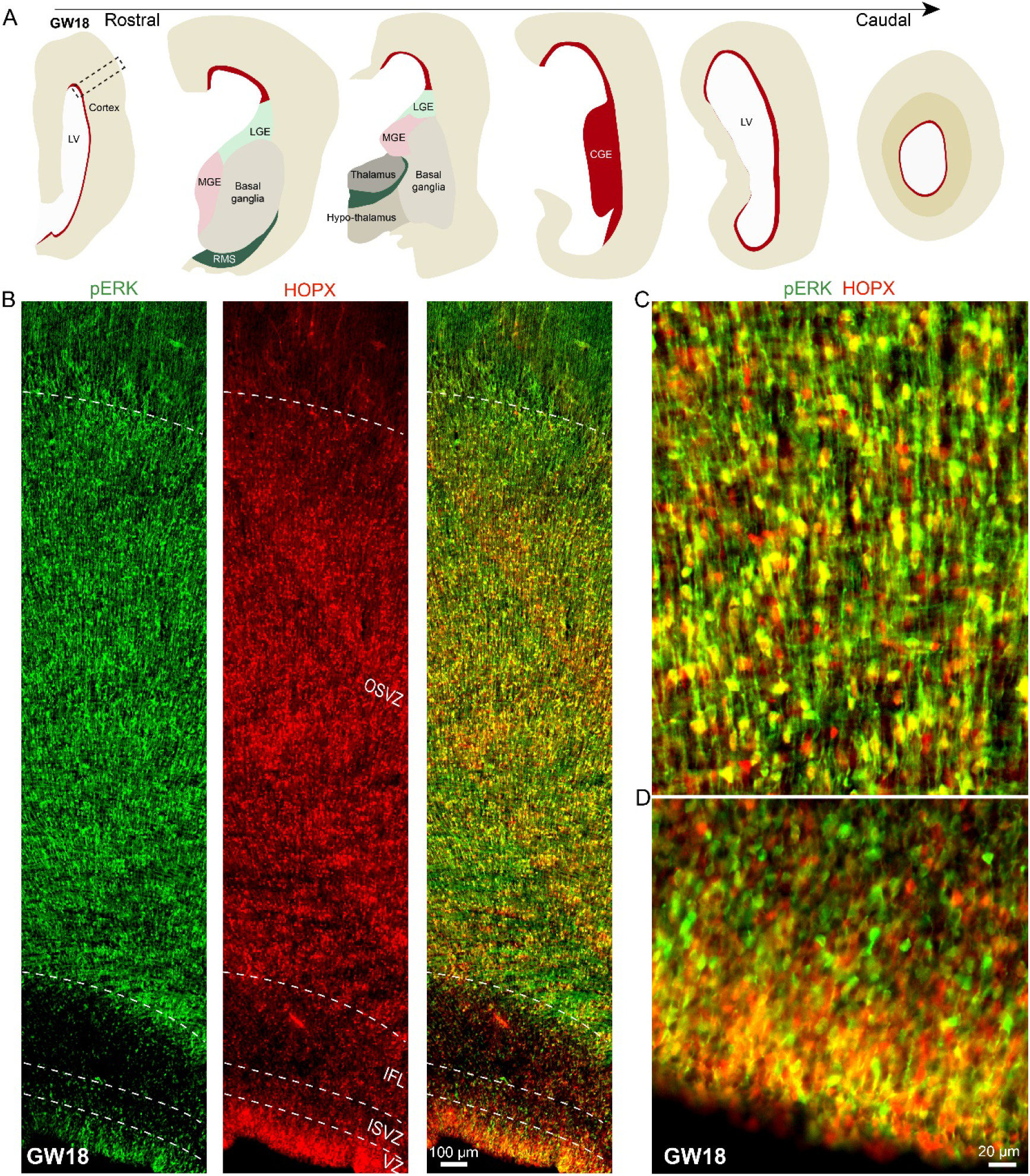
Stronger ERK signaling activity in human cortical oRG cells than tRG cells. **(A)** The diagram showing brain sections spanning the rostral-caudal extent of the human telencephalon at GW18. CGE, caudal ganglionic eminence; MGE, medial ganglionic eminence; RMS, rostral migratory stream. (B) The coronal section through the rostral telencephalon (the outlined area in A) at GW18 double immunostained with pERK and HOPX. ISVZ, inner subventricular zone, IFL, inner fiber layer. (C, D) Higher magnification images showing that pERK and HOPX immunoreactivity is stronger in the cortical OSVZ (C) than cortical VZ (D).

**Figure 7.**
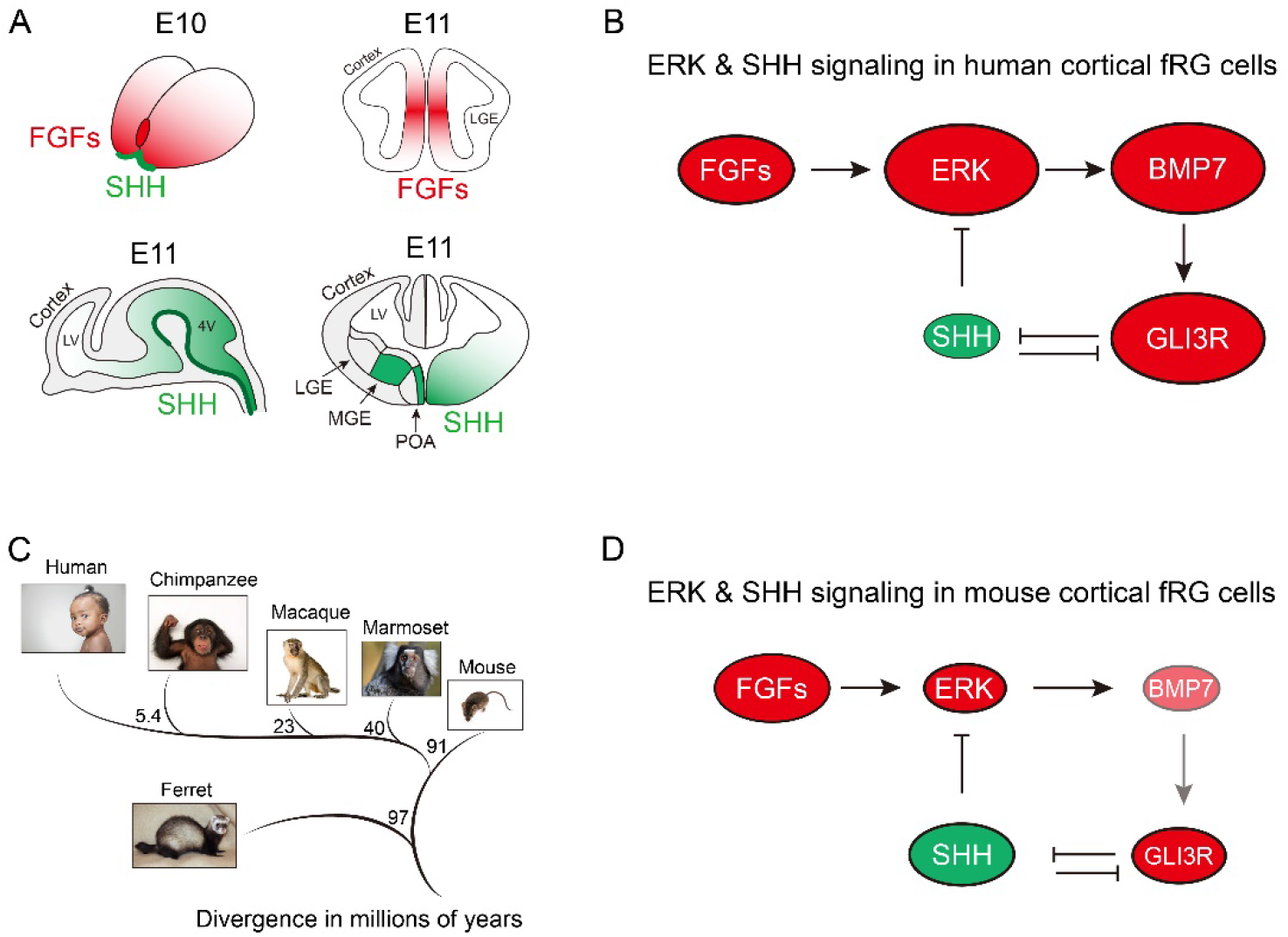
The principle of human and mouse cortical development and evolution. **(A)** During early telencephalon patterning and development, FGFs are expressed in the rostral patterning center, whereas SHH is expressed in the POA and MGE-derived neurons. In general, FGF-ERK signaling exhibits in a rostral^high^- caudal^low^ gradient, whereas SHH-SMO signaling exhibits in a ventral^high^-dorsal^low^ and caudal^high^-rostral^low^ gradient. 4V, fourth ventricle; POA, preoptic area. (B) ERK signaling drives the expansion and evolution of the human cortex. At the beginning of cortical neurogenesis in the most recent ancestor to all mammals, it is assumed that there is already a subset of cortical fRG cells that express relatively higher levels of pERK. Elevated ERK signaling in these cortical fRG cells promotes *BMP7* expression, which increases GLI3R generation and represses SHH signaling. A decrease in SHH signaling in cortical fRG cells further enhances ERK signaling. Therefore, ERK-BMP7-GLI3R signaling pathway in cortical fRG cells participates in a positive feedback loop, which expands the cortical fRG cell pool, increases the length of the cortical neurogenic period, and thus drives cortical development, expansion and evolution. (C) The mouse is lissencephalic but originated from a larger and gyrencephalic ancestor, as gyrencephalic ferrets separated from the human phylogenetic tree before mice and rats. (D) SHH signaling drives mouse cortical evolutionary dwarfism. During mouse cortical development and evolution, smaller brains with the smaller lateral ventricle results in a relatively higher levels of SHH signaling in the cortex, which antagonizes ERK signaling. Relatively weak ERK signaling fails to induce *Bmp7* expression in the dorsal cortical RG cells. Therefore, mouse cortical neurogenesis is mainly protected by GLI3R, but not BMP7, resulting in a shortened period of cortical neurogenesis (for example, from more than 130 days in humans to about 7 days in mice), which is associated with a greatly reduced the number of cortical neurons and cortical size.

Taken together, by integrating analyses of published mouse, ferret, monkey and human cortical scRNA-Seq datasets with our immunostaining and mRNA in situ hybridization experiments, we provide strong evidence that ERK signaling is elevated in cortical RG cells during mammalian development and evolution.

## DISCUSSION

Here, we provide evidence that increased FGF-ERK signaling drives the evolutionary expansion of the mammalian cerebral cortex. There are 5 main findings in this study. (1) FGF-ERK signaling promotes the self-renew and expansion of cortical neuroepithelial and RG cells, which increases the size of cortical neural stem cell pool. (2) FGF-ERK signaling induces *Bmp7* expression in cortical RG cells, which increases the length of the neurogenic period. (3) BMP7 promotes GLI3R formation through the canonical SMAD pathway, which inhibits SHH signaling. (4) SHH signaling antagonizes ERK-BMP7 signaling in cortical RG cells, which promotes cortical gliogenesis. (5) ERK signaling is elevated in mammalian cortical RG cells during development and evolution. Therefore, ERK-BMP7-SHH signaling, which mutually inhibit each other, coordinately regulate cortical development and evolution.

On the basis of these results, we wish to suggest a principle of mammalian cortical development, expansion and evolution (Figure 7). Notably, extensive similarities in expression patterns of FGFs in the anterior signaling domain of the hemichordate Saccoglossus kowalevskii and vertebrates provide compelling evidence that FGF-ERK signaling is an ancient genetic regulatory program that drives brain (neural) development and evolution ^85^.

### ERK signaling drives the expansion of the human cortex

The anterior neural ridge is essential for induction of the neural plate and telencephalon in vertebrates, and this process in general is promoted by FGF-ERK signaling ^1, 70, 86^. After neural tube closure, the dorsal and ventral telencephalon give rise to pallial (cortical and hippocampal) and subpallial structures, respectively. We propose that FGF-ERK signaling continually plays a crucial role in driving mammalian cortical development, expansion and evolution (Figure 7A, B). The most recent ancestor to all mammals, i.e., 100 million years ago, is assumed to have already been in some degree gyrencephalic ^7, 37^, and is assumed to have a subset of fRG cells in the developing dorsal cortex that express relatively higher levels of pERK. Elevated ERK signaling in cortical fRG cells induces *BMP7* expression, which promotes GLI3R production and represses SHH signaling ^6^. A decrease in SHH signaling in cortical fRG cells further enhances ERK signaling. Therefore, ERK- BMP7-GLI3R signaling in cortical fRG cells participates in a positive feedback loop (Figure 7B), which increases the size of cortical RG cell pool, and restrains but allows neurogenesis, while inhibiting gliogenesis, and thus increases the length of the neurogenic period. In the human developing cortex, this ERK-BMP7-GLI3R positive feedback in fRG cells most likely starts at the beginning of neurogenesis (GW7), and ends at the beginning of gliogenesis (GW16). We hypothesize that SHH concentration in cerebrospinal fluid (CSF) is reduced in the enlarged lateral ventricle of big brained mammals during cortical development and evolution, which could contribute to a further decrease in SHH signaling and an increase ERK-BMP7-GLI3R signaling in cortical fRG cells.

In the absence of SHH signaling, GLI3FL (acts as a GLI3 activator) protein is processed to GLI3R that represses SHH signaling. This process depends on adenylyl cyclase-mediated cAMP-dependent protein kinase A (PKA) ^87, 88^. Around midterm gestation, human cortical fRG cells have CXCL12 and CXCR4 expression ^6, 25^, whereas mouse cortical RG cells express extremely low levels of CXCL12 and CXCR4 ^89^. CXCR4 is a GPCR that activates Gαi protein and inhibits cAMP-PKA signaling. Thus, the autocrine activation of CXCL12/CXCR4 in human fRG cells decreases GLI3R production ^90^, increases GLI3FL production and enhances SHH signaling, thereby interrupting the ERK-BMP7-GLI3R positive feedback loop in cortical fRG cells. Previous studies have shown that CXCL12/CXCR4 also activates PI3K/AKT/mTOR and YAP-CCN1/2 signaling ^91, 92^. We propose that SHH signaling (including CXCL12/CXCR4 signaling) combined with ERK signaling are the major forces that drive human cortical *BMP7*-expressing fRG cells to give rise to tRG and oRG cells around GW16 ^6^, and ERK-BMP7 signaling is crucial for generating and maintaining the self-renewing oRG stem cell identity in the OSVZ, although this hypothesis is under investigation.

tRG cells are believed to inherit the apical domain from fRG cells, and tRG cell primary cilia contact the CSF in the lateral ventricle, receiving CSF SHH molecules ^72, 93, 94^. tRG cells also express higher levels of CXCL12/CXCR4 ^6^, which results in expressing higher levels of GLI3FL, and lower levels of GLI3R and *BMP7*. This promotes EGFR expression in tRG cells, as GLI3R and BMP7 strongly inhibit EGFR expression ^6^. The onset of EGFR expression in tRG cells is a strong signal for the switch of neurogenesis-to-gliogenesis ^6, 25, 48, 95, 96^. In contrast, oRG cells do not express CXCL12, but express CXCL14 and CXCR4 ^6^, and CXCL14/CXCR4 likely exerts an inhibitory effect on the CXCL12/CXCR4 signaling pathway ^97^. Therefore, ERK-BMP7-GLI3R expression in oRG cells in the cortical OSVZ continues to participate in a positive feedback loop, which makes oRG cells neurogenic from GW17 to at least GW26, continually producing upper layer PyNs ^6, 25^. The oRG cells do not express EGFR and do not give rise to OPCs ^6, 25^. On the other hand, EGFR-expressing primed tRG cells in the cortical VZ are mainly gliogenic, generating basal multipotent intermediate progenitor cells (bMIPCs) that express EGFR, ASCL1 and OLIG2 ^25^. bMIPCs then undergo several rounds of mitosis and generate most of the cortical oligodendrocytes, astrocytes, and a subpopulation of olfactory bulb interneurons ^25, 48^. This two-germinal-zone system in the human developing cortex ensures that the onset of tRG gliogenesis normally occurs in the VZ, as glial cells have multiple indispensable functions in the developing cortex ^98^, while oRGs in the OSVZ continue to produce neurons ^6, 25^, which significantly increases the length of the neurogenic period, a process that is critical for producing the largest number of cortical PyNs in mammalian kingdom ^2, 7, 36^. Furthermore, human-specific genes with preferential expression in cortical progenitors have also been implicated in cortical expansion and folding ^18, 99–106^.

### SHH signaling drives phyletic dwarfism of the lissencephalic mouse cortex

The lissencephalic mouse is believed to have originated from a larger and gyrencephalic ancestor (Figure 7C) ^7, 37, 38^. This is known as evolutionary dwarfism (phyletic dwarfism), a process in which large animals tend to evolve smaller bodies, followed by the simplification of cortical gyrification with a relatively smaller brain ^37^. Smaller brains with the smaller lateral ventricle may result in higher concentrations of SHH that bath the cortical VZ, and which antagonizes ERK signaling during development and evolution (Figure 7D). Although ERK signaling is gradually elevated in parts of the mouse cortical progenitor domains ^40, 51^, it still fails to induce *Bmp7* expression in the dorsal cortical fRG cells. However, when cortical SHH-SMO signaling is blocked in *hGFAP-Cre; Smo^F/F^* mice, elevated ERK signaling is able to induce *Bmp7* expression in a subset of cortical fRG cells ^6^. Therefore, during mouse cortical development, *Bmp7* expression is restricted to the medial cortex ^6^, in which FGF8/17/18-ERK signaling is stronger than the dorsal cortex ^39, 40, 51^. Thus, mouse cortical neurogenesis is mainly protected by GLI3R, but not BMP7. Relatively lower ERK signaling reduces the size of cortical RG cell pool, and also reduces the length of the neurogenic period as lack of BMP7 protection. After only 7 days neurogenesis, mouse cortical gliogenesis occurs from the lateral cortex to the medial cortex around E17.0 ^48^, whereas human cortical RG cells generate PyNs for 130 days ^6^. This is the main reason that mouse cortex only has ∼13.7 million neurons, whereas human cortex has ∼16.3 billion neurons ^32, 33, 35^.

### Multiple types of genetic evidence support our proposed principle of cortical expansion and evolution

First, the normal development and expansion of the human cortex provide the strongest evidence supporting our conclusion that it is ERK signaling that drives evolutionary expansion of the mammalian cerebral cortex. The human frontal cortex, the center of complex thinking, planning and decision-making, is most highly developed and expanded brain structure in mammal evolution ^5, 13^. FGFs are released at the rostral patterning center, followed by subsequent diffusion of these FGFs, which generates a rostral^high^-caudal^low^ gradient of FGF-ERK signaling in the cortex. This is thought to control the size of the rostral telencephalon ^1, 28, 30, 39, 49, 70, 107^. Here, we identified that ERK-BMP7 signaling pathway plays a crucial role in expanding the size of RG cell pool and increasing the length of neurogenic period ^6^. Furthermore, ERK-BMP7 signaling is elevated during cortical development and evolution; this results in significant expansion of the human cortex, especially the frontal lobe.

Secondly, the abnormal development and expansion of the human temporal and occipital cortex in Thanatophoric Dysplasia (TD) patients provide another strong genetic evidence that supports our conclusion. TD is a lethal form of short-limbed dwarfism caused by abnormal mutations of the *FGFR3*, which lead to constitutive activation of FGFR3 tyrosine kinase activity ^56, 108, 109^. *FGFR3* gain of function mutation in TD patients significantly strengthens ERK signaling ^110, 111^, which may further enhance *BMP7* expression. *FGFR3* is expressed in a high caudomedial-low rostrolateral gradient in the human ^112^, ferret ^113^ and mouse cortical neuroepithelial and RG cells ^50, 52, 114^. Therefore, constitutive activation of the mutant FGFR3 protein (independent of the ligand of FGFs), inducing higher level of ERK-BMP7 signaling in the caudal cortex, causes the significant expansion of temporal and occipital lobe in TD patients ^56, 109^. We propose that mutant *FGFR3* may also enhance ERK-BMP7 signaling in chondrocytes that antagonizes Indian hedgehog signaling through promoting GLI3R formation, which inhibits their proliferation, enhances differentiation, promotes bone formation and fusion of ossification centers, resulting in skeletal dwarfism. Notably, the TD mouse provides an excellent model for the human TD, as they also exhibit severe dwarfism, perinatal lethality and the expansion of the cortex ^108, 110, 115–119^, further indicating that FGF-ERK signaling supports cortical expansion.

Third, in humans, loss of function mutation in *NR2F1* gene cause Bosch-Boonstra-Schaaf optic atrophy syndrome (BBSOAS), a rare neurodevelopmental disorder with vision impairment and intellectual disability. Previous studies have shown reproducible polymicrogyria-like malformations in the parietal and occipital cortex in young BBSOAS patients with de novo mutation of *NR2F1* ^120–122^; this is probably due to the elevation of ERK signaling, as NR2F1 inhibits pERK expression in cortical RG cells ^40, 123^. This hypothesis is further supported by the expansion of posterior cortex in *Nr2f1* mutant mouse embryos ^120^.

Finally, because BMP signaling also functions to maintain stem cells in a quiescent state (Li and Clevers, 2010), we suggest that BMP7- and GLI3R-expressing RG cells in the mouse medial cortex (similar to human cortical fRG and oRG cells), when they receive strong cell proliferation signals, are able to sustain their self-renew, maintain neurogenesis, and inhibit gliogenesis for a longer time than that of lateral cortical RG cells ^6^. Therefore, cortical folding is always observed in the medial cortex in those transgenic mouse models for studying cortical expansion, including *hGFAP-Cre; Rosa^SmoM2^* ^124^, *Emx1-Cre; Gli3 ^F/F^* ^125^, *Emx1-Cre; Cep83 ^F/F^* ^126^, and *hGFAP-Cre; Pik3ca^H1047R^* (conditional activating mutations of *Pik3ca*) ^127^ mouse lines, further highlighting that ERK-BMP7-GLI3R signaling plays a similar role in driving mouse cortical development and evolution.

### Materials and Methods

All procedures involving animals, including ferrets and mice, were carried out according to guidelines described in the Guide for the Care and Use of Laboratory Animals, and were approved by Institutes of Brain Science, Shanghai Medical College, Fudan University.

### Animals

Ten transgenic and knockin mouse lines were used in this study (Figure S1A). *Emx1-Cre* ^64^ (JAX no. 005628), *hGFAP-Cre* ^46^ (JAX no. 004600), *Nes-Cre* ^63^ (JAX no. 003771), *Rosa^Fgf8^* ^45, 128^, *Rosa^SmoM2^* ^129^ (JAX no. 005130), *Rosa^MEK1DD^* ^60^ (JAX no. 012352), *Smo* flox ^130^ (JAX no. 004526), and *Smad4* flox ^131^ (Jax no. 017462) mice were described previously. *Map2k1 (exon 2) flox* and *Map2k2 (exon 4-9)* flox mice were purchased from GemPharmatech, Nanjing (Figure S1A), China. *Rosa^Fgf8^*mice were generously provided by Professor Yanding Zhang at Fujian Normal University. The day of the vaginal plug detection was designated as Embryonic day 0.5 (E0.5). The day of birth was designated as P0. The genders of the embryonic mice were not determined and both male and female postnatal mice were used. One pregnant ferret (Mustela putorius furo) was purchased from Wuxi Sangosho Biotechnology Co., Ltd, Wuxi, China, and we used E33 ferret embryos in this study for immunohistochemistical and *BMP7* mRNA in situ hybridization.

### Plasmid Construction

*pCAG-ires-GFP* plasmid was from Addgene (Addgene #11150). Mouse *Bmp7* cDNA and *sFGFR3c* cDNA ^49, 75^ were cloned and inserted into *pCAG-GFP* vector to construct *pCAG-Bmp7-ires-GFP* and *pCAG-sFGFR3c-ires-GFP* overexpression plasmids.

### In Utero Electroporation (IUE)

Overexpression of plasmids was performed in the cortex using IUE at E14.0 or E15.0. About 0.5 μl of 1-2 μg/μl plasmid solution with 0.05% Fast Green (Sigma) was injected into the lateral ventricle of the embryos using a cable-drawn glass micropipette. Five electrical pulses (duration: 50 ms) were applied across the uterus to E14.0 embryos at 33 V and E15.0 embryos at 35 V. The interval between pulses is 950ms. Electroporation was performed using a pair of 7 mm platinum electrodes (BTX, Tweezerrode 45-0488, Harvard Apparatus) connected to an electrocleaner (BTX, ECM830).

### Fixation and sectioning of the brain tissue

Embryos were harvested from deeply anesthetized pregnant mice. Each embryo was separated from the placenta, the brain was dissected out, and then fixed in 4% diethylpyrocarbonate and paraformaldehyde (DEPC-PFA) overnight. Postnatal mice were deeply anesthetized and perfused intracardially with 4% paraformaldehyde (PFA). One pregnant ferret at E33 was anesthetized and the brains were dissected out. All brains were fixed overnight in 4% PFA at 4°C and cryoprotected in 30% sucrose for at least 24 hours, embedded with O.C.T. (Sakura Finetek) in ethanol slush with dry ice. Mouse and ferret brains in this study were sectioned in a coronal plane at 20 μm. The GW18 human cortical sections were 60 μm, and were used in our previous studies ^25, 132, 133^.

### Immunohistochemistry

All immunohistochemical stains in this study were performed on 20 μm coronal cryostat sections of mouse and ferret embryos and 60 μm coronal cryostat sections of GW18 human brain sections as previously described ^25, 48^. Sections were rinsed with TBS (0.01 M Tris–HCL + 0.9% NaCl, pH = 7.4) for 10 min, incubated in 0.5% Triton-X-100 in TBS for 30 min at room temperature (RT), and then incubated with block solution (5% donkey serum + 0.5% Triton-X-100, pH = 7.2) in TBS for 30 min at RT. For double immunostaining, primary antibodies from different species were incubated simultaneously followed by secondary antibodies. Primary antibodies were diluted in donkey serum block solution and incubated overnight at 4°C, incubated for an additional 30 min at RT, and rinsed three times with TBS. Sections were then incubated with secondary antibodies (1:600, all from Jackson ImmunoResearch) for 2 hours at RT, rinsed three times with TBS for 10 min, incubated with 4’,6-diamidino-2-phenylindole (DAPI, 1:5000, Sigma) diluted in TBS for 5 min, and then finally rinsed three more times with TBS. Primary antibodies used in this study include: goat anti-EGFR (1:1000, R&D System, BAF1280), rabbit anti-OLIG2 (1:500, Millipore, AB9610), rabbit anti-pSMAD1/5/9 (1:100, Cell signaling Technology, 13820S), guinea pig anti-EOMES (1:500, Asis Biofarm, OB-PGP022); rat anti-EOMES (1:500, Thermo Fisher, 12-4875-82), rabbit anti-ASCL1 (1:1,000, Cosmo Bio, SKT01-003), rabbit anti-ID3 (1:5000, Biocheck Inc, BCH-4/17-3), rabbit anti-HOPX (1:500, Proteintech, 11419-1-AP), mouse anti-HOPX (1:500, Santa Cruz Biotechnology, sc-398703), goat anti-SP8 (1/2000, Santa Cruz Biotechnology, sc-104661), rabbit anti PAX6 (1/1000, MBL International, PD022), rabbit anti-ETV5 (1/500, Proteintech, 13011-1-AP), rabbit anti-ERK1/2 (MAPK, 1:400, Cell Signaling Technology, #4370) and chicken anti-GFP (1:3000, Aves labs, GFP-1020). Secondary antibodies against the appropriate species were incubated for 2 h at RT (all from The Jackson Laboratory, 1:500). Omission of primary antibodies eliminated staining. All sections were counterstained with 4#,6-diamidino-2-phenylindole (DAPI) (Sigma, 400 ng/mL, 2 min).

### Mouse and ferret *Bmp7* mRNA in situ hybridization

Mouse and ferret *Bmp7* mRNA in situ RNA hybridization experiment was performed using digoxigenin riboprobes on 20 μm cryostat sections. The sections were first postfixed in 4% PFA for 20 minutes. mRNA in situ hybridization was processed as previous described ^73^. Riboprobes were made from mouse cDNAs amplified by PCR using the following primers: *Bmp7*-F: GGGCCAGAACTGAGTAAAGGAC; *Bmp7*-R: GAAGCTCATGACCATGTCGG; or from ferret cDNAs: ferret *Bmp7-F*: TGATCATGGGCTGTGAAGTCTCA; *Bmp7-R*: CCTTACCTCAAGGAGCTGATAG.

### Bulk RNA-Seq

The whole cortex from E14.5 control mice (n = 5), from E14.5 *hGFAP-Cre; Rosa^Fgf8^* mice (n = 5), and from E14.5 *hGFAP-Cre; Rosa^MEK1DD^* mice (n = 4) (Figure S3C) was dissected. Total RNA was purified with a mini RNA isolation kit (ZymoGenetics). RNA-seq was performed as recommended by the manufacturer (Illumina). Levels of gene expression were reported in fragments per kilobase of exon per million fragments mapped (FPKM) ^134^. A gene was considered to be expressed when it had FPKM > 1. For a gene to be called as differentially expressed, it required p < 0.05. Bulk RNA-Seq data from this experiment were deposited in the Gene Expression Omnibus (GEO) under the accession number GSE240381.

### Cortical single cell dissociation

The whole mouse cortex at E14.5 was dissected out under a microscope for scRNA-Seq analysis and bulk RNA-Seq analysis (Figure S1C). FlashTag (CellTrace Yellow, Life Technologies, #C34567, 0.5 μl of 10 mM) ^135, 136^ was injected into the lateral ventricle of littermate control *Smo^F/F^* or *hGFAP-Cre; Smo^F/F^* mice at E17.0. The cortex was collected 24 hours later (E18.0) for cell sorting and scRNA-Seq analysis. Briefly, mouse embryos were dissected out and immediately submerged in fresh ice-cold Hanks balanced salt solution (Gibco 12175-095). The cortex was then cut into pieces and dissociated into a single-cell suspension using a Papain Cell Dissociation Kit (Miltenyi Biotec, catalog no. 130-092-628) according to the manufacturer’s instructions. FlasgTag labeled single-cells were sorted using a BD FACSAriaII (BD Biosciences).

### Construction of 10X Genomic scRNA-Seq libraries and sequencing

The Chromium droplet-based sequencing platform (10X Genomics) was employed to generate scRNA-Seq libraries, following the manufacturer’s instructions (manual document part number: CG00052 Rev C). The cDNA libraries were purified, quantified using an Agilent 2100 Bioanalyzer, and sequenced on an Illumina Hiseq4000. High-quality sequences (clean reads) of samples were processed using Cell Ranger to obtain quantitative information on gene expression. Cellular quality control thresholds were set at 750-5000 genes and <10% mitochondrial transcripts per cell. After filtering, the number of cells in our dataset were as follows: E14.5 control cortex, 8649 cells, 1390 genes/cell; E14.5 *Emx1-Cre; Map2k1/2-dcko* cortex, 5811 cells, 1713 genes/cell; E14.5 *hGFAP-Cre; Rosa^SmoM2^* cortex, 11868 cells, 2663 genes/cell. These scRNA-Seq data have been deposited in the GEO under the accession number GSE240381. Previously, we have performed scRNA-Seq analyses on E18.0 *Smo^F/F^* (control) cortex, 10524 cells, 1954 genes/cell; *hGFAP-Cre; Smo^F/F^* cortex, 7596 cells, 2104 genes/cell; E39 ferret cortex, 23482 cells, 1759 genes/cell, and these scRNA-Seq data from E18.0 *Smo^F/F^* (control) and *hGFAP-Cre; Smo^F/F^* mice, and E39 ferret cortical scRNA-Seq data have been deposited in the GEO under the accession number GSE221389 ^6^. On the other hand, E15.5 mouse cortex scRNA-Seq data is from ^76^, E78 rhesus monkey visual cortical scRNA-Seq data is from ^77^, and GW14 human cortex scRNA-Seq data is from ^78^, and GW18, GW22, GW23, and GW26 human cortex scRNA-Seq dataset is from ^81^. Raw read counts were processed using the global scale normalization method Log Normalize. The normalized datasets were then combined using the Find Integration anchor and Integration Data functions. Statistically significant principal components identified by resampling tests were retained for unified manifold approximation and projection (UMAP) analysis. The enriched genes in different clusters were identified by Wilcoxon rank sum test (adp value <0.05, |log2FC|>0.25). For cluster annotation, we searched for the most comprehensive and reliable cell type markers through an extensive literature review. All these analyzes were performed in the Seurat package v4.0.6.

### Microscopy and imaging

All images of stained sections in this study were collected by Olympus VS120 Automated Slide Scanner with X20 or X10. Adobe Photoshop software was used to combine multiple visual fields, if needed. Both Adobe Photoshop and Adobe Illustrator were employed to process or adjust images without destroying the original details.

### Quantification and statistical analysis

In this study, all cell counts were performed on the imaged sections collected by Olympus VS120 Automated Slide Scanner with X20. For the quantification of numbers EOMES^+^ and PAX6^+^ cells in the lateral cortex of control and *Emx1-Cre; Map2k1/2-dcko* mice at E17.0, all the measurements were done using sections that match the same anterior–posterior level. At least three matching sections from each brain were used for measurement and compared between mutant and control littermates (n = 3 mice per group). Cells in the cortical sections were counted within 250 µm wide radial segment.

The length of the cortical VZ of the lateral ventricle at E14.5 was determined in DAPI-stained sections by measuring the distance between the medial cortex and lateral cortex. At least 3 sections spanning from anterior to posterior regions were measured and compared between littermate control and mutant brains from 3 pairs of littermates.

The thickness of the cortical VZ and SVZ at E17.0 was also determined in DAPI-stained sections. Quantification of the thickness of the VZ and SVZ was performed along the anterior–posterior axis in littermate control and *Emx1-Cre; Map2k1/2-dcko* mice (n = 3 per group). At least 3 sections from each of the anterior, medial, and posterior telencephalon regions were used.

Results from quantification were analyzed for statistical significance using Student’s paired *t* test. All data are presented as means ± SEM. Significance is stated as follows: p<0.05 (*), p<0.01 (**), p<0.001 (***).

## Supplementary data

The online version contains supplementary data, including 8 Figures and 5 Tables.

## Ethics approval

All procedures and animal care, including ferrets and mice, were approved and performed in accordance with the Fudan University, Shanghai Medical College Laboratory Animal Research Center guidelines.

## Consent to participate

All authors give their consent to participate.

## Consent for publication (include appropriate statements)

All authors give their consent to participate.

## Authors’ contributions

M.S. and Y.G. conducted majority of experiments and analyzed the data. Z.L. designed and interpreted some experiments. L.Y. performed bioinformatics analysis. G.L., Z.X., R.G., Y.Y., contributed to cell counting, partial immunostaining experiments, data analysis, and genotyping. Z.Y. conceived the study, initiated the project, designed and interpreted the experiments, provided the resources, supervised the whole project and wrote the manuscript.

## Data availability

scRNA-Seq data of the E14.5 cortex of control mice (*Map2k1* and *Map2k2* flox), E14.5 cortex of *Emx1-Cre; Map2k1/2-dcko* mice, E14.5 cortex of *hGFAP-Cre; Rosa^SmoM2^* mice, and bulk RNA-Seq data of E14.5 cortex of control mice (n =5), E14.5 cortex of *hGFAP-Cre; Rosa^Fgf8^* mice (n = 5), E14.5 cortex of *hGFAP- Cre; Rosa^MEK1DD^*mice (n = 4), used in this study have been deposited in the Gene Expression Omnibus (GEO) under the accession number GSE240381. The E18.0 *Smo^F/F^*(control) and *hGFAP-Cre; Smo^F/F^* mouse cortex (mainly including RG cells and progenitors), and E39 ferret cortex scRNA-seq data have been deposited in the Gene Expression Omnibus (GEO) under the accession number GSE221389 ^6^. All datasets generated in this study are available upon request.

## Acknowledgments

We thank Professor J. L. Rubenstein for comments, suggestions and edits. We are grateful to Professor Y. Zhang at Fujian Normal University for sharing the *Rosa^FGF8^* mouse.

## Abbreviations

bMIPCs: basal multipotent intermediate progenitor cells
CGE: caudal ganglionic eminence
CSF: cerebrospinal fluid
DEPC: diethylpyrocarbonate
fRG: full span radial glia
GLI3FL: GLI3 full-length
GLI3R: repressor form of GLI3
GPCR: G protein-coupled receptor
GW: gestational week
IFL: inner fiber layer
IPCs: intermediate progenitor cells
ISVZ: inner subventricular zone
IUE: in utero electroporation
LGE: lateral ganglionic eminence
LV: lateral ventricle
MGE: medial ganglionic eminence
oRG: outer radial glia
OSVZ: outer subventricular zone
PFA: paraformaldehyde
PKA: protein kinase A
POA: preoptic area
PyNs: pyramidal neurons
RG: radial glia
RMS: rostral migratory stream
scRNA-Seq: single cell RNA sequence
tRG: truncated radial glia
VZ: ventricular zone.

