## Supplemental Figures for "ERK signaling drives evolutionary expansion of the mammalian cerebral cortex"

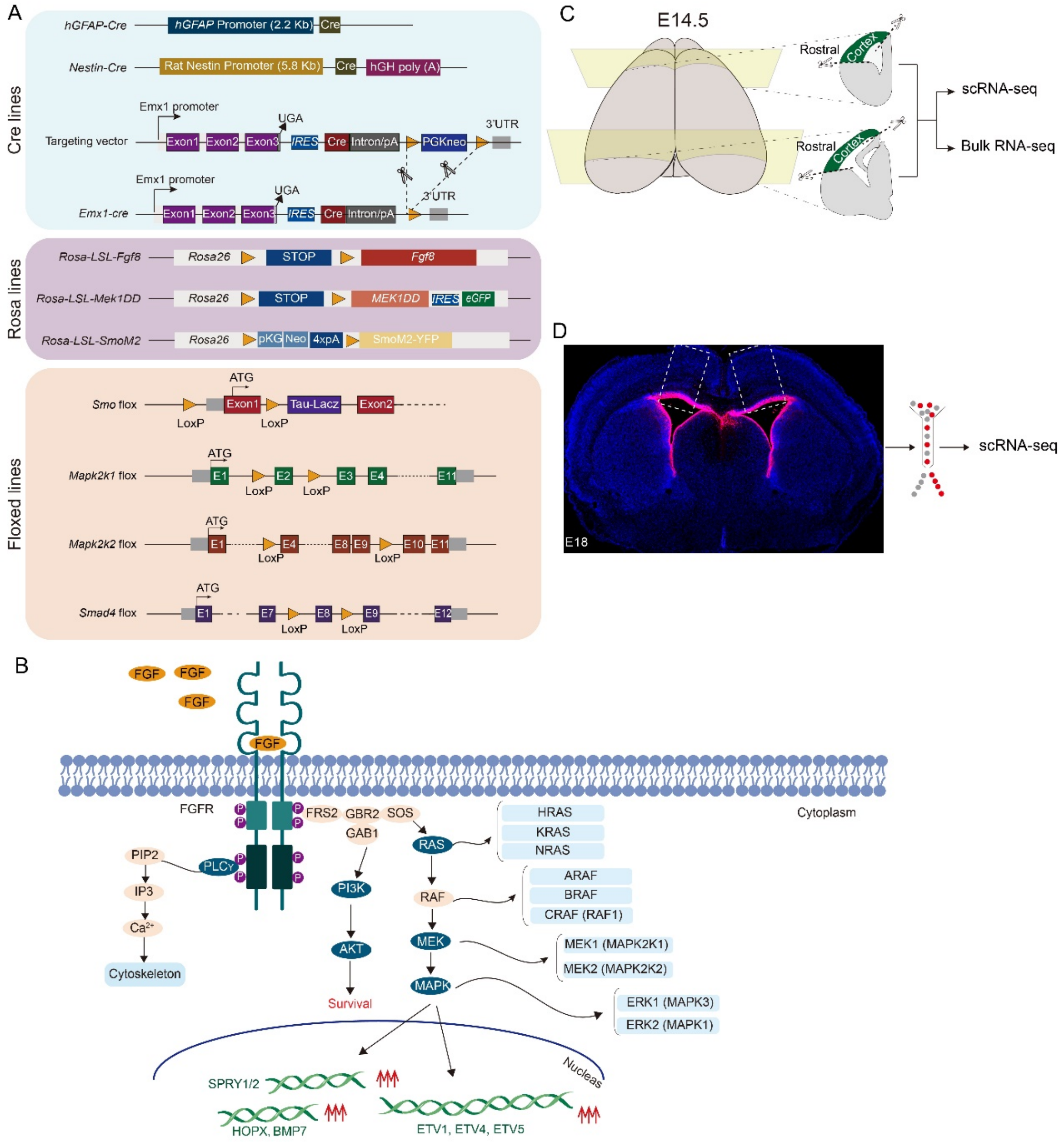


**Figure S1. The strategies and tools for studying FGF-ERK signaling and SHH-SMO signaling in the developing mouse cortex.** (A) The list of transgenic and knockin mouse lines used in this study, including *hGFAP-Cre*, *Emx1-Cre*, *Nes-Cre*, and $Rosa^{FGF8}$, $Rosa^{MEK1DD}$, $Rosa^{SmoM2}$, *Map2k1* flox, *Map2k2* flox, *Smo* flox, *Smad4* flox mice. (B) Schematic illustration of signaling pathways activated downstream of FGF signaling. (C) Schematic of the workflow of bulk RNA-Seq and scRNA-Seq analysis of the mouse cortex at E14.5. The whole cortex was directly dissected from the E14.5 telencephalon. (D) Schematic of the workflow of scRNA-Seq analysis of mouse cortical RG cells at E18.0. FlashTag (CellTrace Yellow) was injected into the lateral ventricle of $Smo^{F/F}$ (littermate control) and *hGFAP-Cre;* $Smo^{F/F}$ mice at E17.0. The cortex was collected at E18.0 for cell sorting and scRNA-Seq analysis.

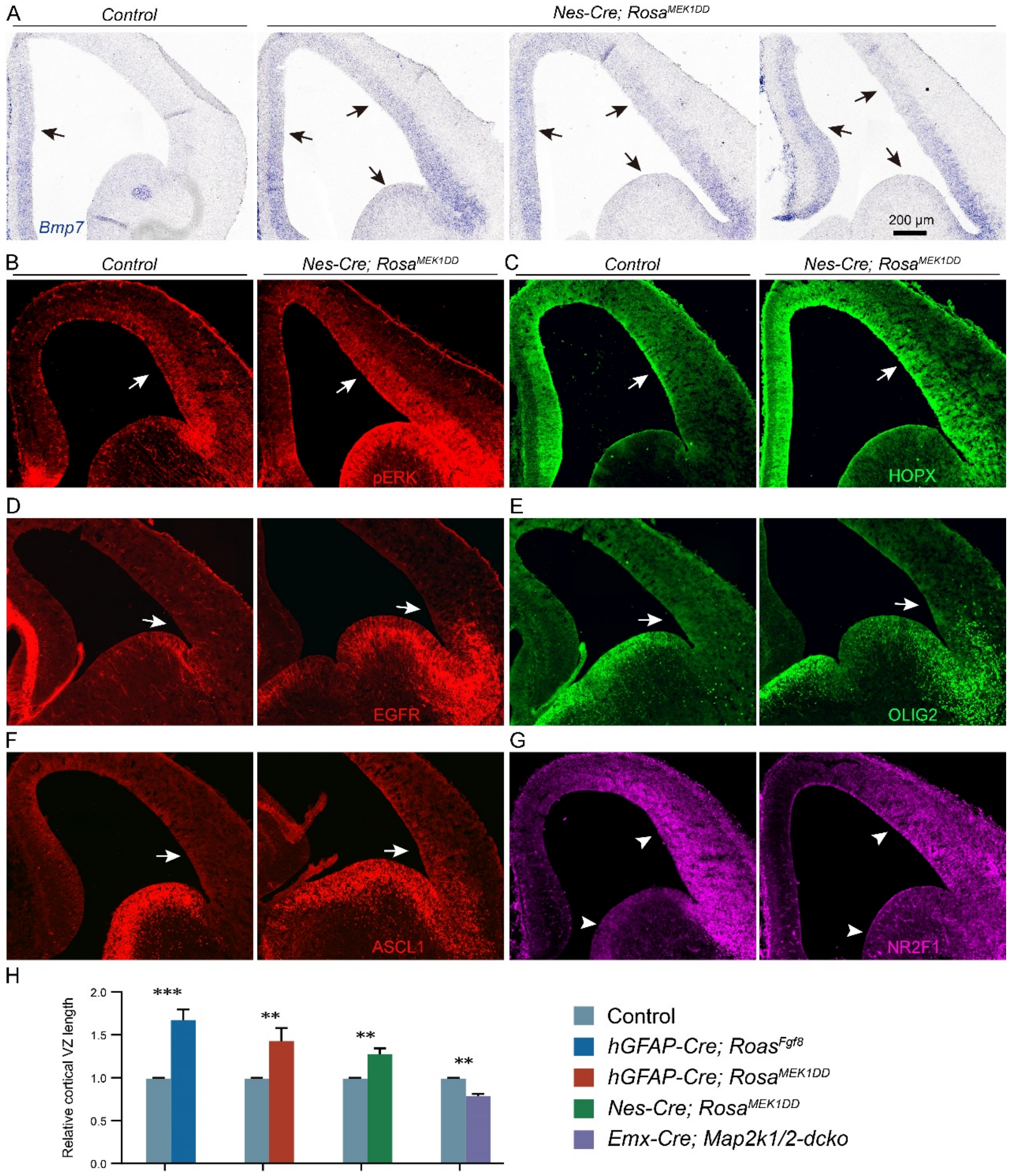


**Figure S2. Sustained activity of ERK signaling in the cortex and LGE of *Nes-Cre; Rosa*$^{MEK1DD}$ mouse at E14.5.** (A-F) Upregulation of *Bmp7*, pERK, HOPX, EGFR, OLIG2 and ASCL1 in the cortex and LGE of *Nes-Cre; Rosa*$^{MEK1DD}$ mice (arrows). (G) NR2F1 expression was downregulated in the cortex and LGE (arrowheads). (H) Quantification of the cortical VZ length in littermate control and transgenic mice.

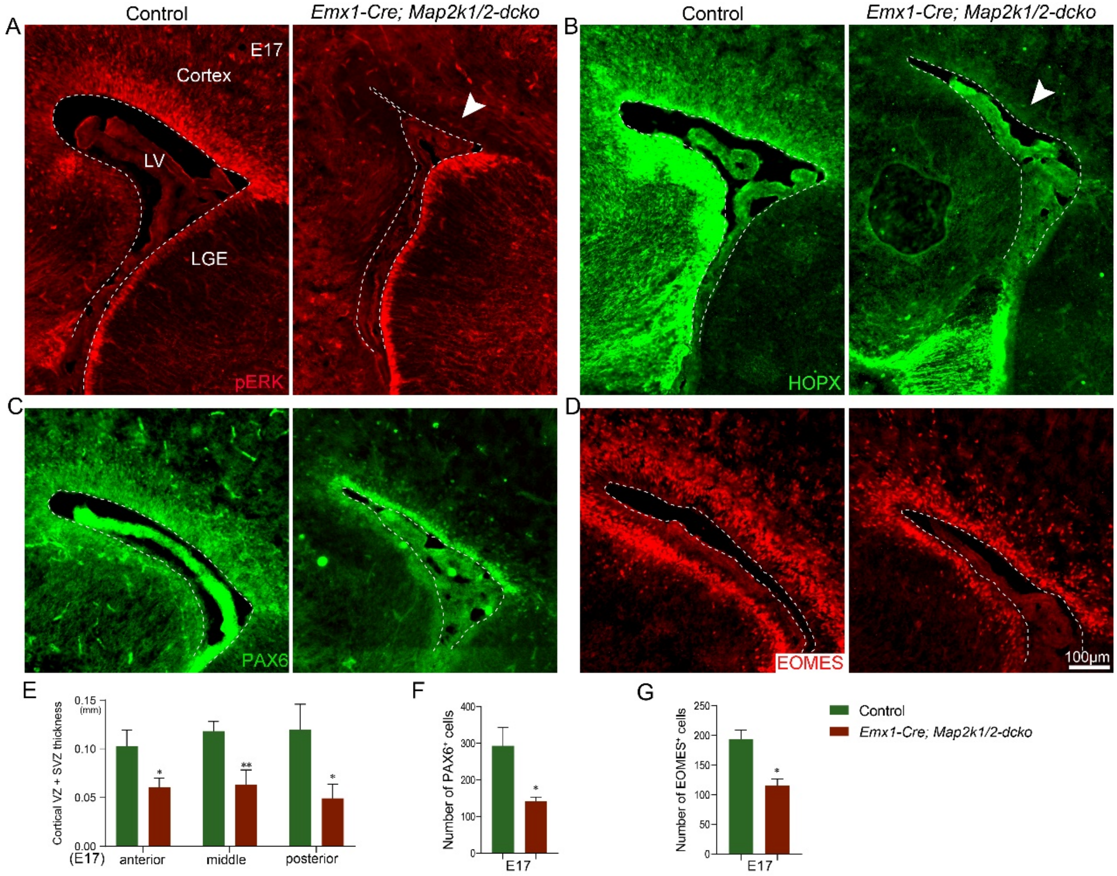


**Figure S3. Loss of ERK signaling leads to a thinner cortical VZ and SVZ.** (A, B) Expression of pERK and HOPX was completely lost in the cortex of *Emx1-Cre; Map2k1/2-dcko* mice at E17.0 (arrowheads). (C, D) PAX6 and EOMES immunostaining of coronal brain sections from a littermate control and *Emx1-Cre; Map2k1/2-dcko* mutant. (E) Quantification of cortical VZ and SVZ thickness based on PAX6 immunostaining. Coronal sections along the anterior–posterior axis of the telencephalon reveal a significantly decrease in the thickness of the cortical VZ and SVZ at E17.0. (F, G) Numbers of PAX6$^{+}$ and EOMES$^{+}$ cells in the cortex were significantly reduced in the *Emx1-Cre; Map2k1/2-dcko* mutant at E17.0, due to loss of cortical RG cells.

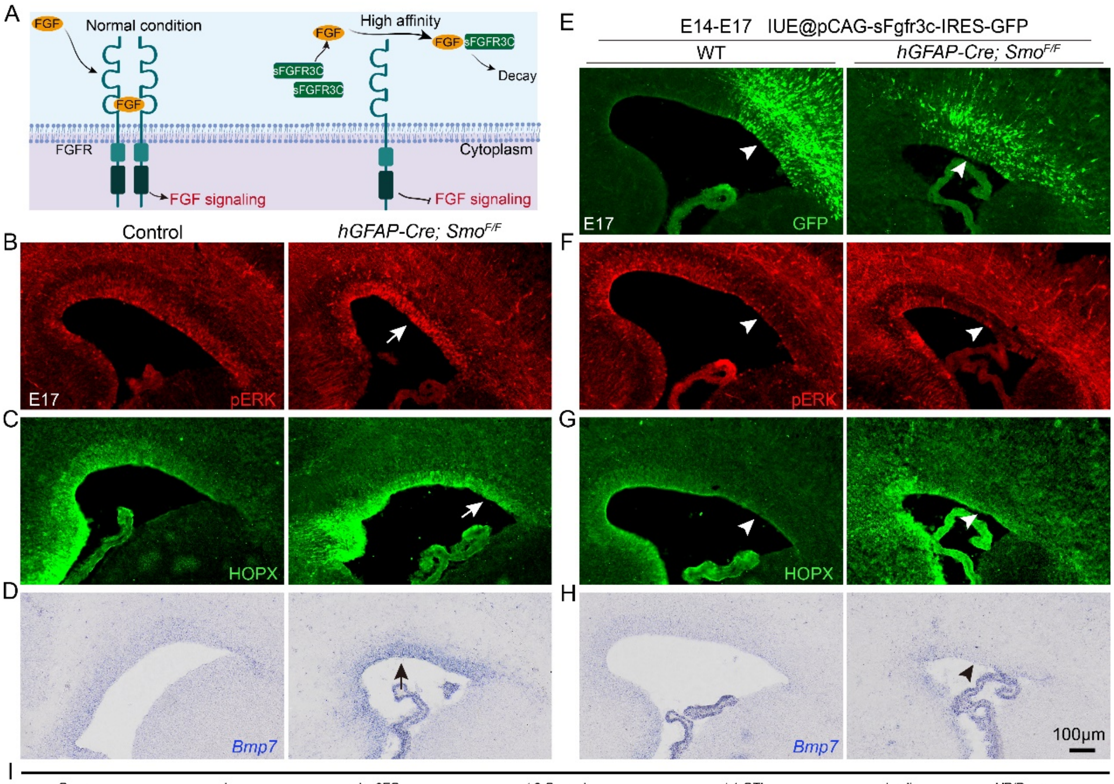


I

| Gene | p_val | avg_log2FC | pct.2-Smo-cko | pct.1-CTL | p_val_adj | UP/Down |
|---|---|---|---|---|---|---|
| *Aldoc* | 4.56E-33 | -0.456 | 0.991 | 0.982 | 7.83E-29 | UP |
| *Bmp7* | 2.58E-05 | -0.055 | 0.141 | 0.074 | 4.42E-01 | UP |
| *Dio2* | 4.40E-48 | -0.653 | 0.407 | 0.100 | 7.55E-44 | UP |
| *Etv1* | 1.72E-03 | -0.094 | 0.485 | 0.364 | 1.00E+00 | UP |
| *Etv4* | 5.01E-19 | -0.327 | 0.331 | 0.141 | 8.59E-15 | UP |
| *Etv5* | 2.07E-12 | -0.213 | 0.476 | 0.273 | 3.56E-08 | UP |
| *Fgfr2* | 1.60E-03 | -0.089 | 0.389 | 0.288 | 1.00E+00 | UP |
| *Gfap* | 6.93E-08 | -0.207 | 0.430 | 0.281 | 1.19E-03 | UP |
| *Hopx* | 5.76E-24 | -0.516 | 0.911 | 0.772 | 9.89E-20 | UP |
| *Id3* | 3.81E-53 | -1.300 | 0.698 | 0.351 | 6.53E-49 | UP |
| *Ncam1* | 7.21E-38 | -0.626 | 0.828 | 0.548 | 1.24E-33 | UP |
| *Nog* | 1.93E-10 | -0.184 | 0.274 | 0.139 | 3.32E-06 | UP |
| *Tnc* | 2.02E-10 | -0.315 | 0.948 | 0.868 | 3.48E-06 | UP |
| *Birc5* | 8.32E-05 | 0.289 | 0.060 | 0.109 | 1.00E+00 | DOWN |
| *Ccnd1* | 6.31E-22 | 0.535 | 0.056 | 0.204 | 1.08E-17 | DOWN |
| *Cd9* | 1.17E-91 | 1.009 | 0.807 | 0.951 | 2.00E-87 | DOWN |
| *Gas1* | 9.68E-25 | 0.717 | 0.586 | 0.685 | 1.66E-20 | DOWN |
| *Slc1a3* | 3.88E-38 | 0.573 | 0.915 | 0.955 | 6.65E-34 | DOWN |
| *Sparcl1* | 9.06E-83 | 1.425 | 0.048 | 0.447 | 1.56E-78 | DOWN |

**Figure S4. Loss of SHH-SMO signaling in the mouse cortex increases ERK signaling in cortical RG cells.** (**A)** Schematic representation of sFGFR3c-mediated inhibition of FGF-ERK signaling in the brain. (**B-D**) Loss of *Smo* function resulted in upregulation of pERK, HOPX, and *Bmp7* (arrows). (**E-H**) Following *sFGFR3c* IUE in the E14.0 cortex that blocked FGF-ERK signaling, the expression of pERK, HOPX, and *Bmp7* was greatly downregulated in the E17.0 cortex of both *Smo*$^{F/F}$ (littermate control) and *hGFAP-Cre; Smo*$^{F/F}$ mice (arrowheads). (I) scRNA-Seq analysis revealed differentially expressed genes (DEG) in E18.0 cortical RG cells of *hGFAP-Cre; Smo*$^{F/F}$ mice (Smo-cko) relatively to *Smo*$^{F/F}$ littermate controls (CTL). Expression of ERK signaling response genes, *Aldoc, Bmp7, Dio2, Etv1, Etv4, Etv5, Fgfr2, Gfap, Hopx, Nog,* and *Tnc* was significantly upregulated.

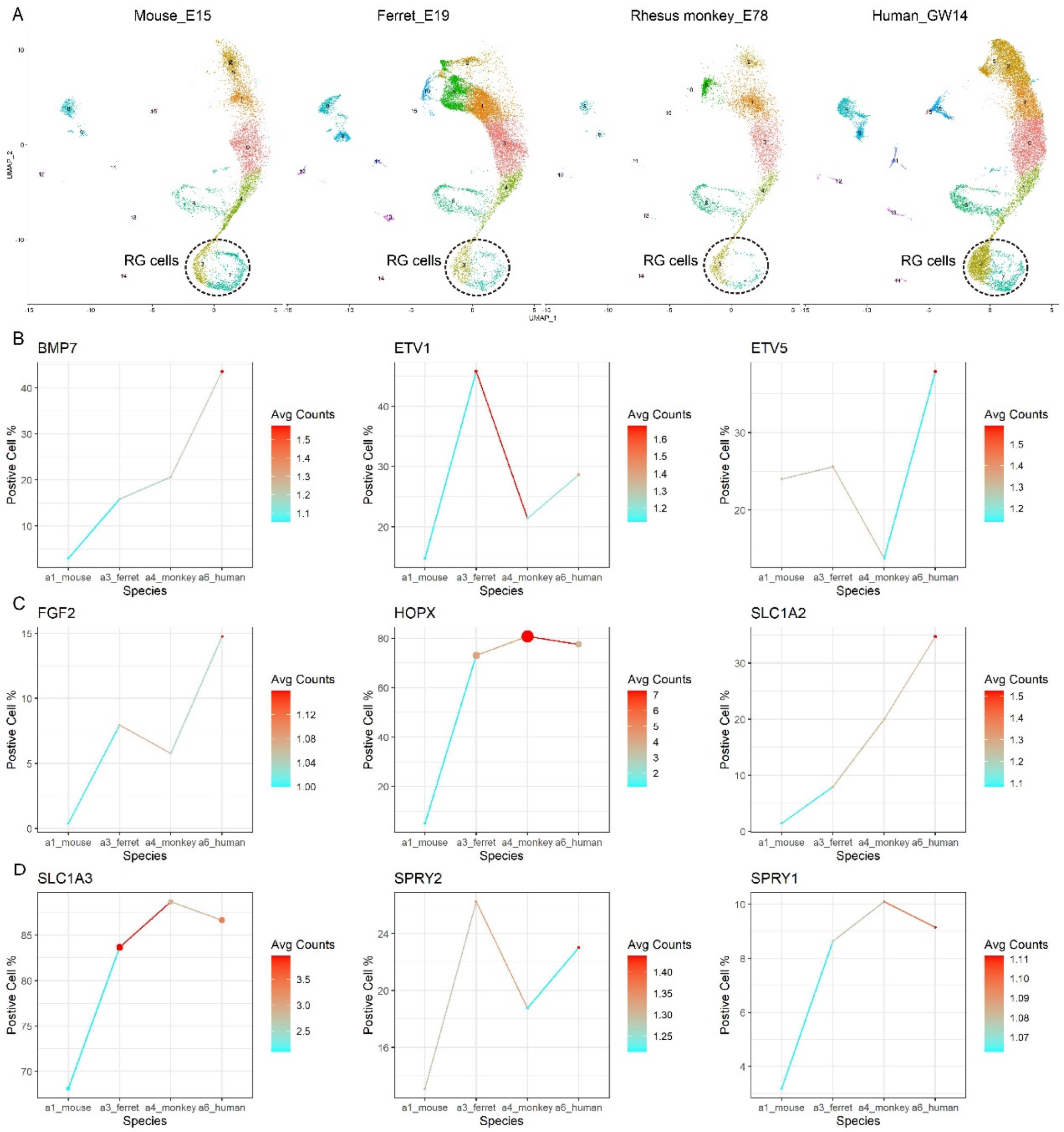


**Figure S5. An evolutionary increase in ERK signaling in mammalian cortical fRG cells. (A)** scRNA-Seq analyses of the developing cortex of mouse, ferret, rhesus monkey and human at the neurogenic stage. Cross-species analyses of transcriptomic signatures of cortical cells from E15.5 mouse cortex, E39 ferret cortex, E78 rhesus monkey visual cortex, and GW14 human cortex. (**B**) In general, there is an increase in expression of *BMP7*, *ETV1, ETV5, FGF2, HOPX, SLC1A2, SLC1A3, SPRY1* and *SPRY2* in cortical fRG cells during evolution.

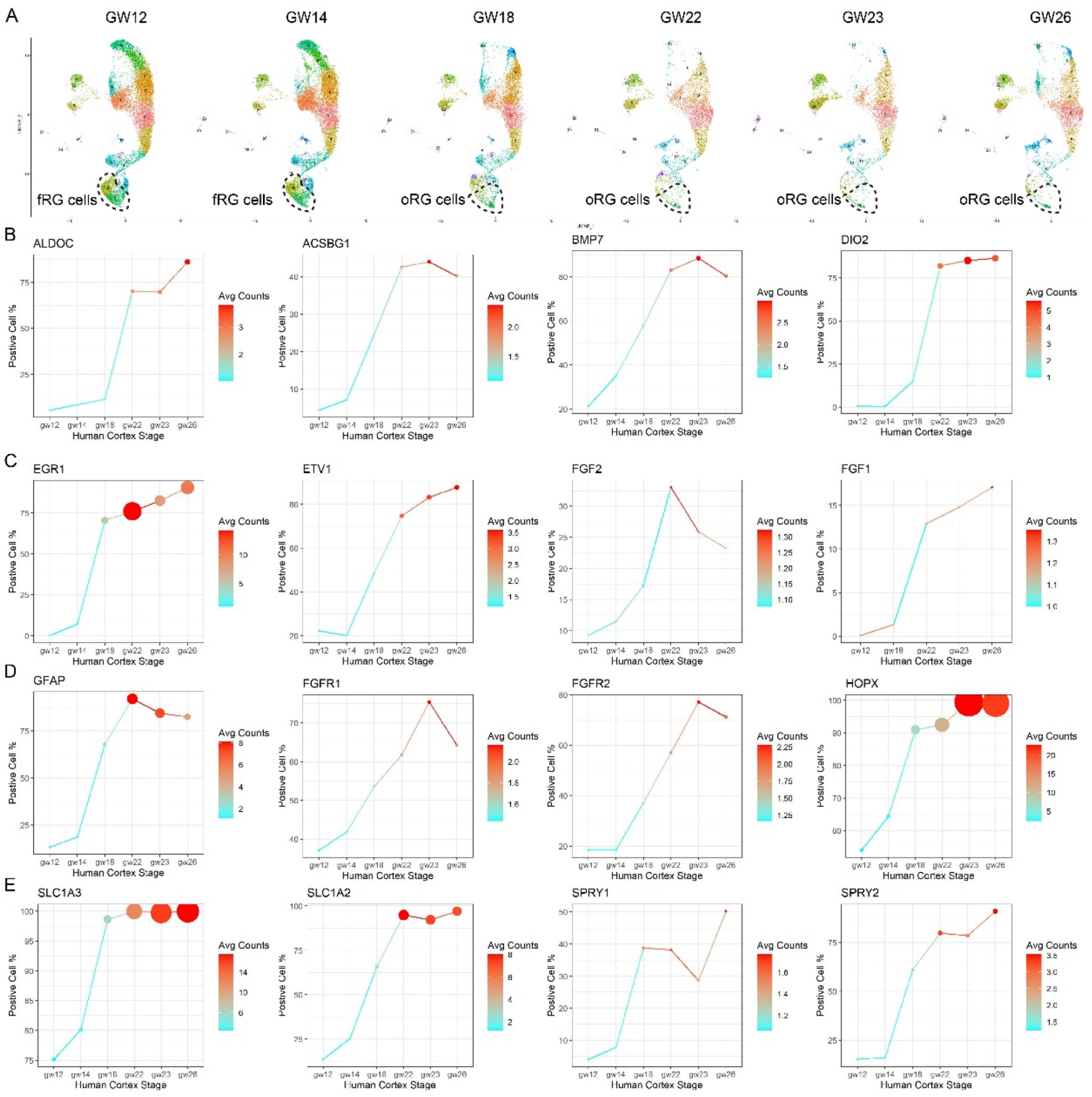


**Figure S6. ERK signaling activity is elevated in human cortical fRG and oRG cells with increasing gestational age.** (A) scRNA-Seq analyses of the developing human cortex at GW12, GW14, GW18, GW22, GW23, and GW26. (B-E) Expression of ERK signaling response genes was increased in cortical fRG and oRG cells with increasing gestational age.

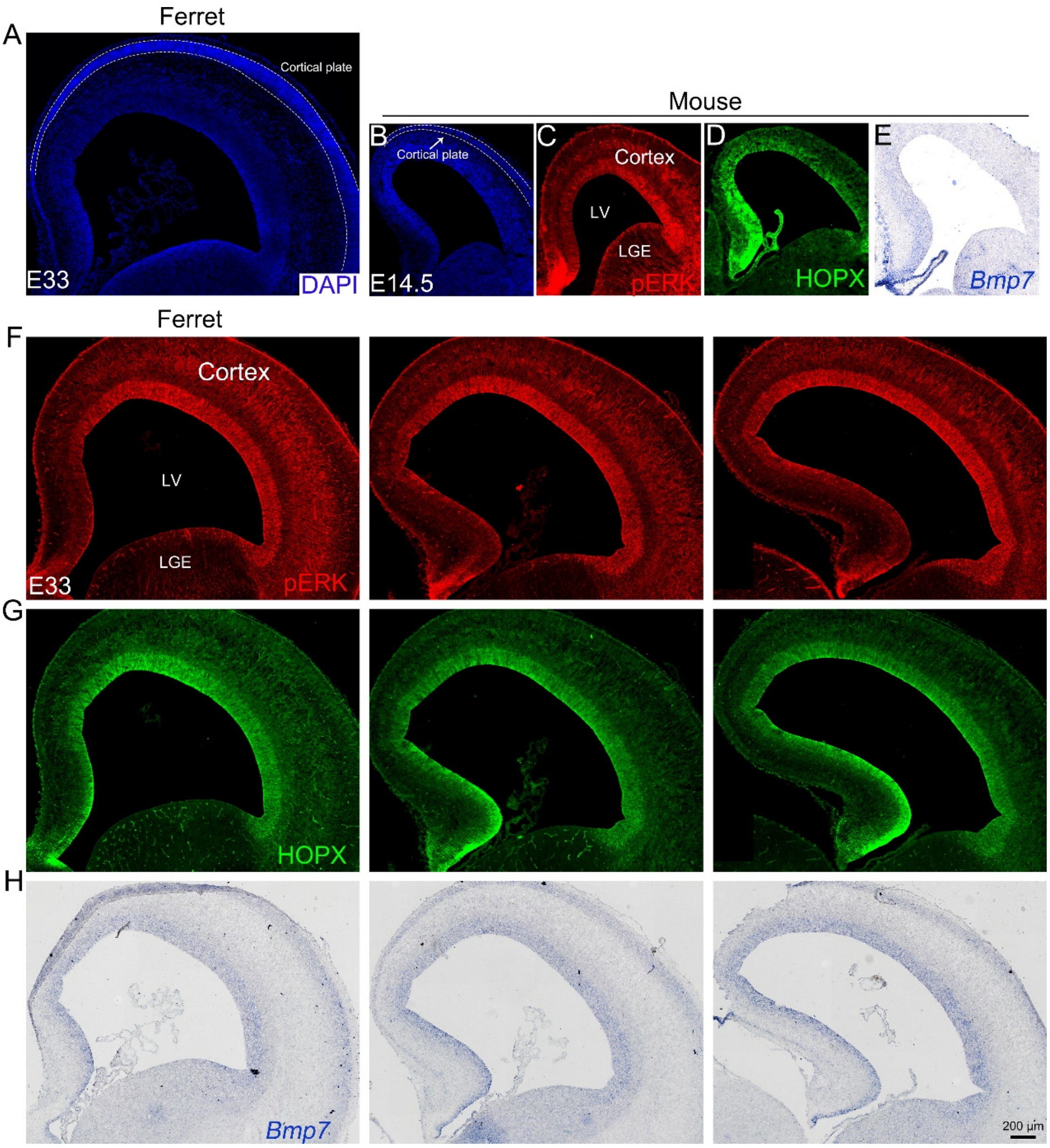


**Figure S7. Stronger ERK signaling in cortical fRG cells in ferrets than mice. (**A, B**)** The coronal cortical sections of E33 ferret and E14.5 mouse embryos were stained with DAPI. The cortical plate is newly formed. (C-H) Expression of pERK, HOPX and *BMP7* in the E33 ferret cortical VZ appeared to be increased compared with the E14.5 mouse cortical VZ.

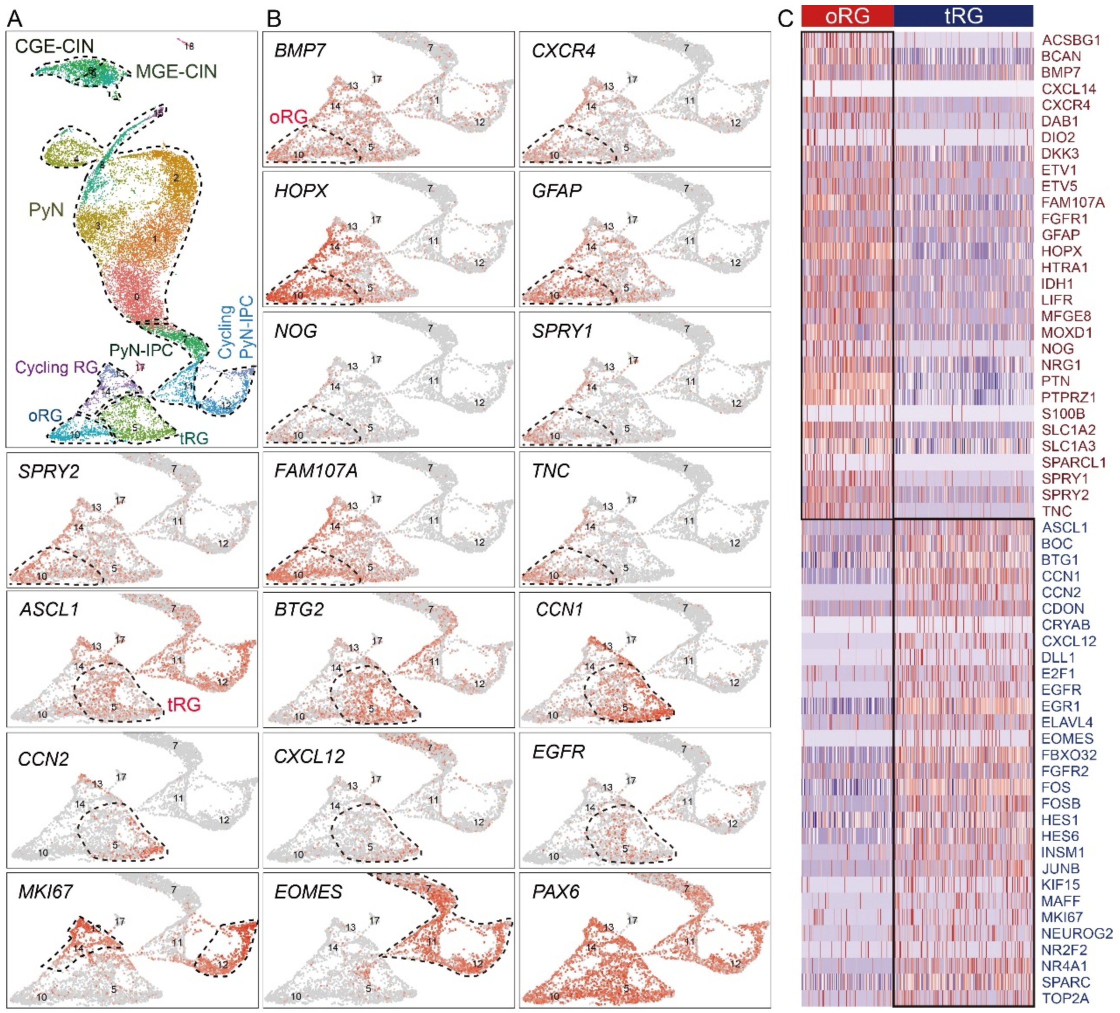


**Figure S8. Stronger ERK signaling in GW18 human cortical oRG cells than tRG cells.** (A) scRNA-Seq analyses of the human cortex at GW18. (B) Expression plots of example differentially expressed genes between cortical oRG and tRG cells. (C) Heat map of gene expression for select genes in oRG and tRG cells, including ERK response genes.

**Table S1**. Bulk RNA-Seq analysis of gene expression in the E14.5 cortex of *hGFAP-Cre; Rosa*$^{Fgf8}$ mice (n = 5) relatively to littermate controls (n = 5).

**Table S2**. RNA-Seq analysis of gene expression in the E14.5 cortex of *hGFAP-Cre; Rosa*$^{MEK1DD}$ mice (n = 5) relatively to controls (n = 4).

**Table S3.** scRNA-Seq analysis of gene expression in E14.5 cortical RG cells in *Emx1-Cre; Map2k1/2-dcko* mice relatively to littermate controls.

**Table S4.** scRNA-Seq analysis of gene expression in E14.5 cortical RG cells in *hGFAP-Cre; Rosa*$^{SmoM2}$ mice relatively to controls.

**Table S5.** scRNA-Seq analysis of GW18 human cortex; full list of upregulation and downregulation genes in oRG relatively to tRG cells (cluster 10 vs. cluster 5).
